# Expression attenuation as a mechanism of robustness to gene duplication in protein complexes

**DOI:** 10.1101/2020.07.09.195990

**Authors:** Diana Ascencio, Guillaume Diss, Isabelle Gagnon-Arsenault, Alexandre K Dubé, Alexander DeLuna, Christian R. Landry

## Abstract

Gene duplication is ubiquitous and a major driver of phenotypic diversity across the tree of life, but its immediate consequences are not fully understood. Deleterious effects would decrease the probability of retention of duplicates and prevent their contribution to long term evolution. One possible detrimental effect of duplication is the perturbation of the stoichiometry of protein complexes. Here, we measured the fitness effects of the duplication of 899 essential genes in the budding yeast using high-resolution competition assays. At least ten percent of genes caused a fitness disadvantage when duplicated. Intriguingly, the duplication of most protein complex subunits had small to non-detectable effects on fitness, with few exceptions. We selected four complexes with subunits that had an impact on fitness when duplicated and measured the impact of individual gene duplications on their protein-protein interactions. We found that very few duplications affect both fitness and interactions. Furthermore, large complexes such as the 26S proteasome are protected from gene duplication by attenuation of protein abundance. Regulatory mechanisms that maintain the stoichiometric balance of protein complexes may protect from the immediate effects of gene duplication. Our results show that a better understanding of protein regulation and assembly in complexes is required for the refinement of current models of gene duplication.

## Introduction

Gene duplication and divergence is a primary source of functional innovation and diversity. During the last few decades, the long-term maintenance of gene duplicates through the gain or reciprocal loss of function has been studied extensively (Ohno 1970; Lynch and Force 2000; Qian and Zhang 2008). However, we know relatively little about the immediate impact of duplications, which may have significant consequences on the preservation of paralogs. Genes with adverse effects on fitness upon duplication would have a reduced residence time in populations, thereby limiting their contribution to long term evolution. For instance, even modest changes in gene dosage such as those caused by duplication sometimes produce significant phenotypic effects, both positive and negative (Iskow et al. 2012; Qian and Zhang 2014; Payen et al. 2016; Rice and McLysaght 2017a). This is commonly referred to as dosage-sensitivity. Understanding the immediate impact of duplication is, therefore, of paramount importance.

Several mechanisms have been suggested to explain dosage-sensitivity. These include concentration dependency, promiscuous off-target interactions at high concentration, and dosage imbalance (Rice and McLysaght 2017b). The gene-balance hypothesis predicts that the single-gene duplication of protein complex subunits is harmful, as this can lead to an immediate stoichiometric imbalance with the rest of the subunits (Papp et al. 2003; Birchler and Veitia 2007; Rice and McLysaght 2017b). There is indirect evidence supporting such a prediction: complex subunits are less likely to be retained after small-scale duplication, are often co-expressed at similar levels, and are enriched among genes that reduce fitness when underexpressed (Papp et al. 2003; Gonçalves et al. 2017). For instance, haploinsufficiency, a dominant phenotype in diploid organisms that are heterozygous for a loss-of-function allele, is more common among genes that encode protein complex subunits (Deutschbauer et al. 2005). However, other works have shown that genes that are toxic when overexpressed are not enriched as part of protein complexes (Sopko et al. 2006; Semple et al. 2008). This suggests that overexpression can be deleterious for reasons unrelated to complex stoichiometry.

While gene deletion and overexpression experiments inform us of how cells respond to lowered or increased abundance of proteins, they are fundamentally different from a naturally occurring duplication. For instance, the use of non-native promoters and multicopy plasmids may cause the assayed genes to be overexpressed several-fold and may also alter the timing of expression. In addition, if dosage-fitness relationships are non-linear, results from overexpression cannot be interpolated to gene duplication. For these reasons, and because fitness rather than complex assembly has been assayed in previous experiments (Sopko et al. 2006; Moriya et al. 2012), we do not know what is the impact of duplication on the assembly of protein complexes. Experiments aimed at measuring the fitness benefits of increased gene dosage have been performed (Payen et al. 2016) but have not addressed how such dosage changes impact protein complex formation.

One reason why the overexpression of protein complex subunits can be less detrimental than a reduction of expression is dosage regulation (Semple et al. 2008). The correct dosage of protein subunits appears to be tightly regulated by the cell. For instance, members of multiprotein complexes are produced in precise proportion to their stoichiometry in both bacteria (Li et al. 2014) and eukaryotes (Taggart and Li 2018). In *Saccharomyces cerevisiae*, proportional synthesis is sometimes maintained by tuning the protein synthesis rates for duplicated genes that encode the same subunit (Taggart and Li 2018). Additionally, recent studies revealed that the abundance of members of multiprotein complexes is often attenuated or buffered in aneuploids (strains with extra copies of one or several chromosomes) (Dephoure et al. 2014; Gonçalves et al. 2017; Chen et al. 2019). A study by Dephoure *et. al* (2014) suggested that attenuation may be quite common since nearly 20% of the proteome is attenuated in aneuploids and most attenuated genes (60-76%) are members of multiprotein complexes (Dephoure et al. 2014). In fact, (Chen et al. 2019) found that up to 50% of subunits with imbalanced gene copy numbers (compared with the rest of the complex) may be attenuated to normal protein abundance levels. Since attenuated genes often have mRNA transcript levels and protein synthesis rates proportional to their gene-copy number, the regulation of attenuated genes most often, but not exclusively, occurs posttranslationally (Dephoure et al. 2014; Ishikawa et al. 2017; Taggart and Li 2018). However, aneuploid cells may not be the best models to study the effect of small-scale gene duplications because a large proportion of their genome is duplicated at once. In addition, aneuploid cells often experience systemic effects like proteotoxic stress (Oromendia et al. 2012), which may cause pleiotropic consequences on protein synthesis and degradation rates that are challenging to disentangle from those of the duplication of a single gene. Furthermore, when a complete chromosome is duplicated, more than one subunit -- or the complete set of subunits of a complex -- may be duplicated, which could lessen or even prevent the effects of stoichiometric imbalance. Therefore, the extent and the nature of attenuation after small duplication events are yet to be explored.

In this work, we sought to measure the immediate impact of gene duplication by experimentally simulating gene duplication of essential genes in yeast. We focused on this set because genes that are essential for growth are enriched in protein complexes (Papp et al. 2003). Genes that are essential are also less likely to be duplicated (He and Zhang 2006), which means that additional copies in the genome will not confound the results of duplication. We measured the fitness consequences of individual gene duplication for nearly 900 genes in individual strain competitions. Duplication of protein complex subunits is not more deleterious on average than that of other genes. We therefore also measured changes in protein-protein interactions (PPIs) in response to gene duplication for a subset of essential genes that are part of four large protein complexes. We also estimated the extent to which the expression of proteasome subunits is attenuated as a response to gene duplication. Our results show that even though gene duplication often affects fitness, it has a small effect on the assembly of protein complexes. The apparent robustness of multi-protein complexes to gene duplication is likely to be a consequence of expression attenuation.

## Results

### An important percentage of yeast’s essential genes affect fitness when duplicated

We measured the fitness effects of gene duplication using high-resolution competition assays with fluorescently labeled cells in co-culture (Figure 1A). We individually duplicated 899 essential genes using single-copy centromeric plasmids (pCEN) expressing the genes under their native promoters and UTRs (Ho et al. 2009). In parallel, we generated a distribution of control strains in which we competed a *WT* strain (also used as a reference) with itself. Each of the 192 replicates of this control set is an independent colony from a transformation.

**Figure 1.**
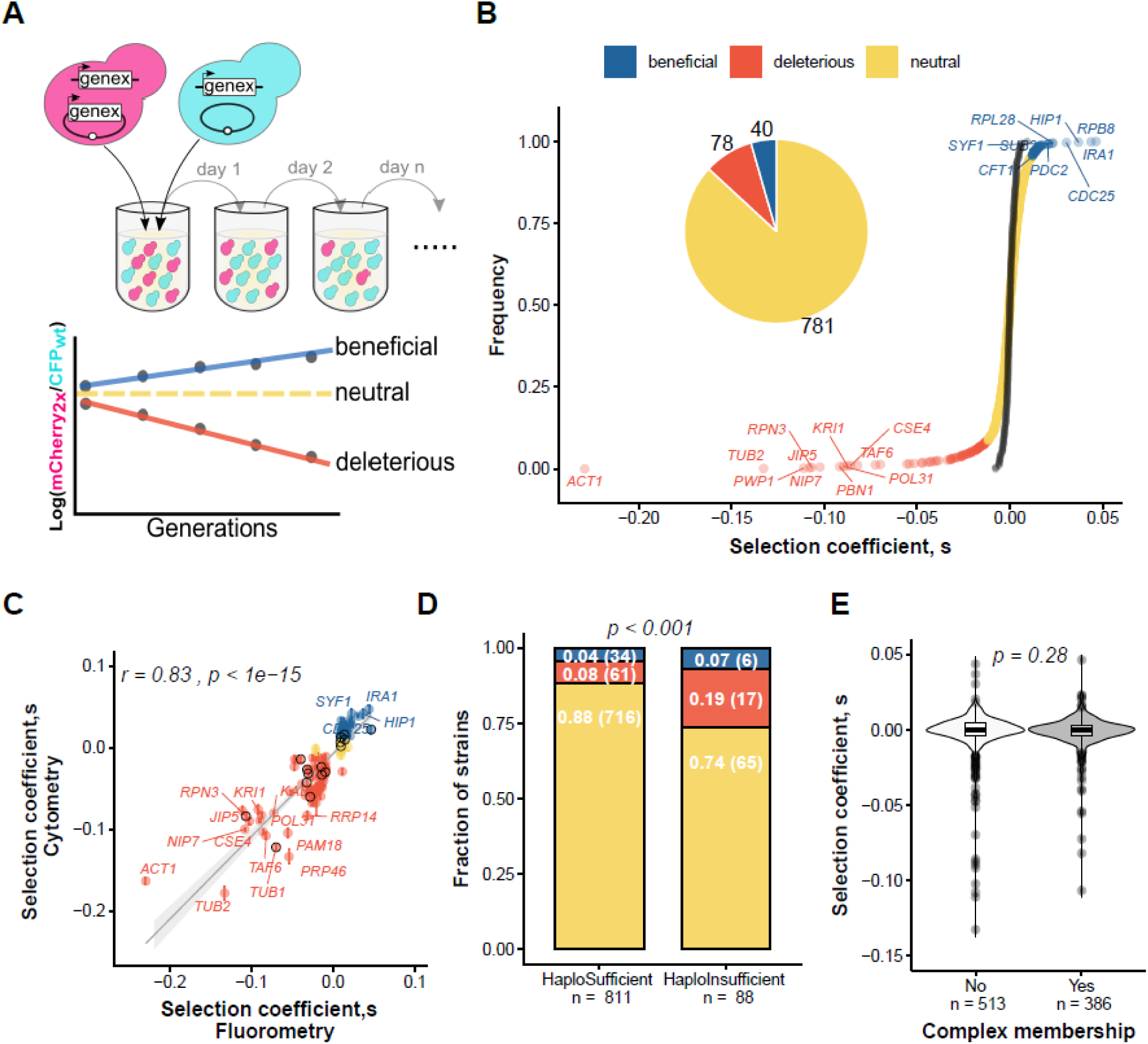
More than 10% of yeast essential genes affect fitness when duplicated. (**A**) Relative fitness was measured using a high-resolution competition assay (DeLuna et al. 2008). We co-cultured a mCherry- tagged strain carrying an extra-copy of an essential gene on a centromeric plasmid with a CFP-tagged reference strain carrying a control plasmid. We followed the ratio of the two populations for up to 28 generations to calculate a slope, which corresponds to the selection coefficient (*s*). (**B**) Cumulative distribution of selection coefficients of all the 899 strains tested (*Supplementary Table S1*). Each dot represents a strain expressing an additional gene copy. The black dots represent the distribution of 192 biological replicates of reference-versus-reference competition. The threshold used for deleterious (in red) or beneficial (in blue) effect is at least 1% (*-4*.*5 > z score > 4*.*5*). (**C**) Selection coefficients for the validation of the 180 genes with significant effects measured by flow cytometry (*Supplementary Table S2*). The labels are for genes with the strongest deleterious and beneficial effects. The bars indicate the standard deviation of three biological replicates. The black circles highlight genes with haploInsufficient phenotypes. Spearman’s correlation coefficient is indicated at the top. (**D**) A comparison of fitness effects among haploinsufficient and haplosufficient genes (Deutschbauer et al. 2005). P-value from a Fisher’s exact-test is shown. The fraction and number of genes are indicated with white letters. (**E**) Selection coefficients of genes that code for proteins that are members of complexes and proteins that are not. On top, we show the *p-value* from a Wilcoxon’s Rank-Sum test.

We chose a conservative threshold of at least 1% percent of fitness effect (*-0*.*01* > *s > 0*.*01*), that corresponds to a |*z-score*| > 4.5. For most genes (86%) duplication has little or no effect on relative fitness. However, around 9% (Figure 1B) of the duplications have moderate to strong deleterious effects (*s < −0*.*01, z-score < −4*.*5*) while 4% have beneficial (*s > 0*.*01, z-score > 4*.*5*) but often modest effects. We validated the top 180 genes (top 180 absolute z-scores) using the same growth conditions but monitoring the two populations by flow cytometry. We generated the strains *de novo* and tested three biological replicates per sample. While less scalable, this approach allows for a more accurate estimation of population ratios since fluorescence is measured at the single-cell level instead of the population level. There is a strong correlation between the two assays (Spearman’s *r = 0*.*88, p < 1e-15*, Figure 1C), giving us good confidence in the accuracy of our large-scale measurements. We validated the vast majority of the effects of the first competition assay (159/180, Figure 1C). We compared our results with two other studies evaluating the relative fitness of strains harboring the same plasmids through pooled assays (Payen et al. 2016; Morrill and Amon 2019). We found a weak but significant correlation between the selection coefficient from these studies and ours (*Supplementary Figure S1*). Although these two previous studies have used a similar pooling approach, they are only weakly correlated with each other (*r = 0*.*15, p<1e-7*) (*Supplementary Figure S1-B*). The weak correlations could come from the different media used (SC-ura versus rich media, or nutrient limitation). Furthermore, pool-assays create a complex and distinct environment compared with pairwise competition assays and may lead to noisier estimates, for instance, strains that are rare in the pools.

The four percent of the duplications have beneficial effects above 1% include genes such as *CDC25* and *IRA1* (Figure 1B-C), which regulate the Ras-cAMP pathway (Broek et al. 1987; Tanaka et al. 1989; Russell et al. 1993). In yeasts, growth and metabolism in response to nutrients, particularly glucose, are regulated to a large degree by this pathway. Since we used glucose as a carbon source, duplication of these two genes may modify the activity of the pathway in a way that increases the growth rate. Adaptation in limited glucose conditions often involves mutations in these pathways (Venkataram et al. 2016). The duplication of some genes that encode central metabolism enzymes were also beneficial. For instance, duplication of *HIP1*, a histidine transporter (Tanaka and Fink 1985) and *PDC2*, a pyruvate decarboxylase (Schmitt and Zimmermann 1982; Velmurugan et al. 1997) may result in increased growth rate through metabolic activity in a similar manner to duplication of genes in the Ras-cAMP pathway.

The presence of beneficial duplications may appear surprising since the yeast lineage has a high rate of duplication (Lynch et al. 2008) and has undergone a whole-genome duplication (Wolfe and Shields 1997; Kellis et al. 2004), which would have provided the mutational input needed to fix any beneficial duplication. One possible reason for the lack of duplication is that the adaptive value of some gene duplications is highly dependent on environmental conditions and can even become deleterious in specific contexts (Kondrashov 2012). Such antagonistic pleiotropy has been observed in the study of aneuploidies (Pavelka, Rancati, Zhu, et al. 2010) and gene deletions (Qian et al. 2012). We, therefore, performed a parallel competition experiment in a condition of salt stress. We find a significant correlation in the selection coefficient between conditions (Spearman’s *r = 0*.*48, p < 1e-15*), and a similar number of deleterious (8.5%) and beneficial (4.6%) effects (*Supplementary Figure S2*). Many of the effects may, therefore, be general while others are condition specific. The overlap in the identity of the deleterious genes between conditions is greater than for advantageous ones, suggesting that beneficial effects are more condition-specific than deleterious ones. To further validate the condition-specificity of the benefit of gene duplication, we measured the relative fitness of five strongly beneficial and three strongly deleterious duplications in five different conditions. We observed antagonistic pleiotropy for some beneficial duplications (*Supplementary Figure S3*). For instance, *IRA1* and *CDC25* were beneficial in the standard condition, osmotic stress and with galactose as carbon source but are strongly deleterious in 6% of ethanol, while having no detectable effect in the presence of caffeine. On the other hand, *ACT1* and *TUB2* have similar deleteriousness across the five conditions tested when duplicated. The mechanisms by which an increase in gene dosage is sometimes beneficial remains to be determined. In the presence of antagonistic pleiotropy, it is possible that expression in any given condition is not optimal but rather represents a tradeoff in terms of adaptation across conditions. Indeed, a study looking at the fitness effects of changes in expression levels of several genes showed that the *WT* expression level in some conditions is often not optimal (Keren et al. 2016).

Deleterious duplications are more frequent and have stronger effects than advantageous ones and include genes such as *TUB1, TUB2* and *ACT1*. The products of these genes are involved in the structural integrity of the cell cytoskeleton and have been previously shown to be highly sensitive to dosage increase (Burke et al. 1989; Katz et al. 1990; Espinet et al. 1995). Our results confirm recent observations that doubling or halving the expression of *TUB1* and *TUB2* is enough to reduce fitness (Keren et al. 2016). These observations suggest that genes that are deleterious upon duplication could also be haploinsufficient. We indeed find that 19% of haploinsufficient genes (Deutschbauer *et al*. (2005) tested here produce deleterious effects when duplicated (17/88, Figure 1D), which is more than the 8% expected by chance (61/811; *Pearson’s χ*^*2*^ *test p < 1e-3*; Figure 1D). Conversely, Qian *et al*. (Qian et al. 2010) suggested that haploinsufficient genes may be beneficial when duplicated. We see a tendency in this direction but it is not significant (6/88 and 34/811; *Pearson’s χ*^*2*^ *test p = 0*.*388;* Figure 1D).

Our interpretation regarding the fitness effects of duplication depends on whether gene expression from a centromeric plasmid approximates the dosage effect of gene duplication, which we expect to commonly be a doubling of dosage (exceptions have been shown, with higher expression than expected (Loehlin and Carroll 2016)). Centromeric plasmids usually segregate as yeast chromosomes and are on average found in one copy per haploid cell (Clarke and Carbon 1980), but it is possible that the number of plasmids varies (Gnügge and Rudolf 2017). Payen *et al*. (Payen et al. 2016) confirmed that in glucose-limiting conditions, the plasmids we used are typically found in only one copy per cell so our results are overall representative of individual gene duplication events. We also examined if the plasmid copy genes are regulated similarly as the genome-encoded copy. We reasoned that if pCEN are systematically present in multiple copies, a protein expressed from a plasmid would result in higher expression than when expressed from the genome in an equivalent genetic background where we have a genomic copy and a plasmid copy. We compared protein abundance of a strain with an endogenous GFP fusion and a copy of the gene on a pCEN plasmid, with the same strain but this time having the GFP fusion expressed from the pCEN plasmid. Only one gene out of five showed higher protein abundance when expressed on the plasmid, and one showed reduced expression (*Supplementary Figure S4-A*). The two genes with different expression levels the differences are rather modest (less than one-fold) as opposed to orders in magnitude that are common in multi-copy plasmids. We also directly measured the abundance of Act1p in the presence of an additional gene copy on a plasmid (pCEN-*ACT1)* by Western-Blot and found only a modest increase in abundance (*Supplementary Figure S4-B*), which is inconsistent with its deleterious effects being driven by several-fold expression changes. These observations suggest that our systematic strategy using pCEN constructions is a good experimental approximation of naturally occurring duplications in terms of dosage.

### The duplication of individual subunits rarely affects PPIs in complexes

The dosage balance hypothesis predicts that both under- and over-expression of a protein complex subunit would cause deleterious phenotypes because dosage perturbation affects the stoichiometric balance between the subunits of the complex and compromises its assembly (Papp et al. 2003; Birchler and Veitia 2007). For instance, some subunits of the RNA polymerase II (Rpb2p) and of the proteasome (Rpn3p) are both haploinsufficient and deleterious when duplicated. This indicates that such proteins are sensitive to changes in gene-dosage in both directions, just as the gene balance hypothesis predicts. However, when we mapped the fitness effects of gene duplication on annotated yeast protein complexes, we found no significant difference compared to genes that are not in complexes (*Wilcoxon rank-sum test p = 0*.*28;* Figure 1E). In fact, both gene categories share a similar percentage of genes with deleterious effects (8% and 9%, respectively). These findings are robust to different *z-score* thresholds (*Supplementary Table S3*). Therefore, the duplication of members of protein complexes is not particularly associated with a decrease in fitness, in apparent contradiction with the dosage balance hypothesis.

Our results suggest that either doubling gene-dosage does not affect the assembly protein complexes, or it affects their assembly but these effects have no particularly strong effects on fitness. Therefore, we next aimed at directly measuring if gene duplication affects protein-protein interactions (PPIs) within complexes *in vivo*. We selected complexes that are sensitive to some but not all gene-dosage changes (the 26S proteasome and the three RNA polymerases, *Supplementary Figures S5*). We measured pairwise PPIs between all pairs of subunits before and after the duplication of each subunit using a Protein-fragment Complementation Assay (PCA) based on the DHFR enzyme (DHFR-PCA, (Tarassov et al. 2008)). The quantitative nature of PCA allows us to estimate a perturbation score (*ps*) as a direct measure of the effects of gene duplication on the PPI network of a complex (Figure 2A).

**Figure 2.**
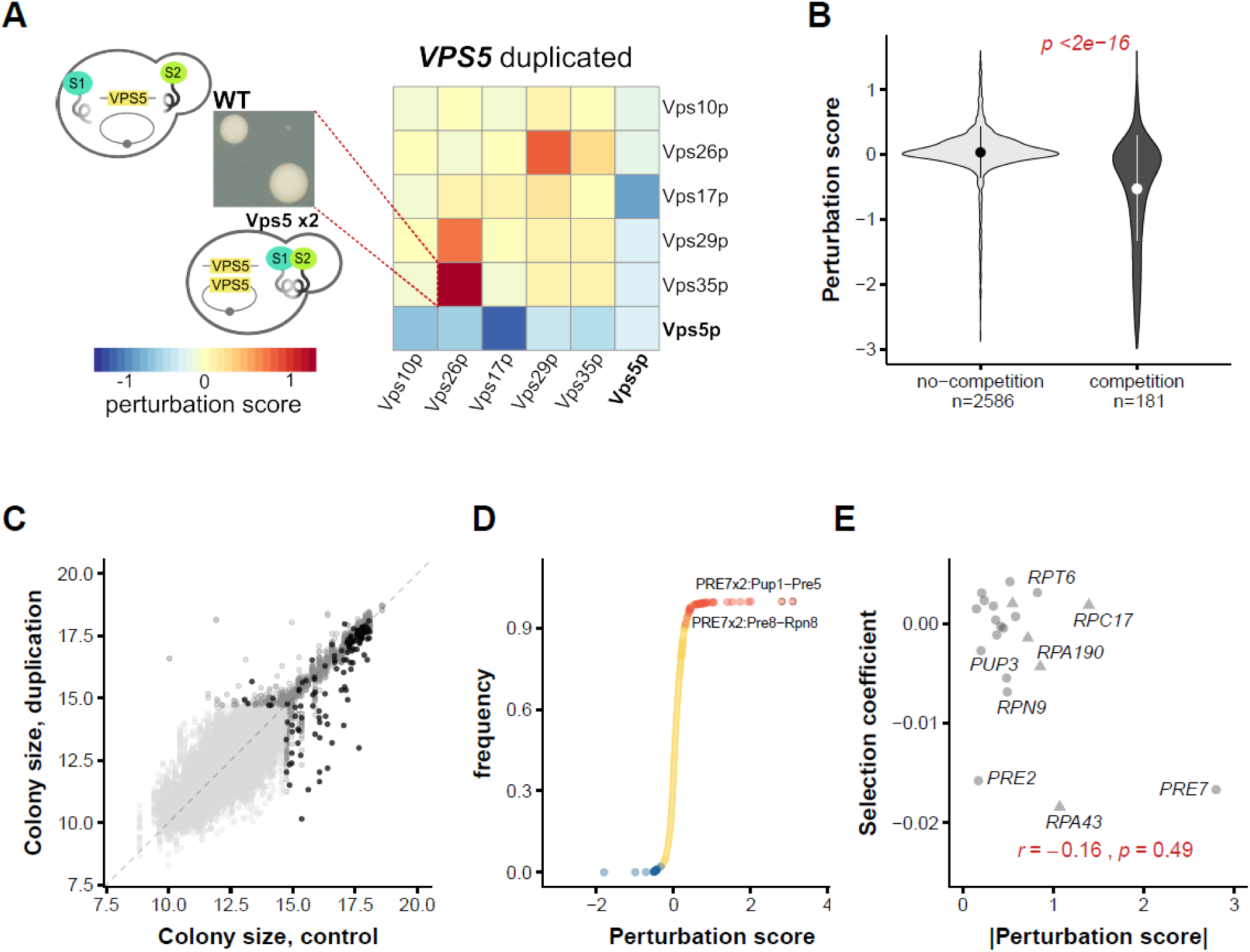
Most duplications of subunits do not affect PPIs in large complexes. (**A**) DHFR PCA-based strategy to measure perturbation of the pairwise physical interactions after a duplication. On the left, in a DHFR-PCA the colony size on selective media (MTX) is correlated with the stability and strength of the physical interaction between the two subunits S1 and S2 (shown in green). The perturbation-score (*ps*) is defined as the colony size difference between the strain carrying a duplication (+pCEN-*VPS5*) and a *WT* (+pCEN) strain. Heatmap indicating *ps* values of the complete retromer PPI network due to the duplication of *VPS5*. (**B**) Distributions of *ps* for interactions with and without a competing subunit. Since the duplicated protein is not tagged with a DHFR fragment, it titrates PPI partners away from the tagged copy, decreasing colony size. We show violin plots of the distributions of all the interactions tested for five complexes (*Supplementary Table S5*). On top, we show results from a Wilcoxon’s Rank-Sum test. (**C**) Colony sizes of all strains carrying the duplication of a subunit compared with their control strain (the empty vector). Colony sizes of diploid strains carrying all tested combinations of preys, baits, and duplications for the proteasome, and the three RNA polymerases (*Supplementary Table S4-5*). Dark gray circles indicate strains above our growth threshold indicative of physical interaction while the light-gray circles are strains below the growth threshold. Black circles indicate interactions with a competing subunit above the threshold. (**D**) Cumulative frequency of *ps* of the proteasome and the three RNA polymerases. All competition effects were excluded. Labeled are the prey-bait combinations that are perturbed by the duplication of *PRE7*. (**E**) Relationship between the selection coefficient and the average *ps* (absolute value) of duplicated subunits on PPIs. Only significant (*FDR of 5%*) and non-competition combinations were used to calculate the averages. Circles represent duplications of proteasome subunits, while triangles represent subunits of any of the three RNA polymerases. In red we show Spearman’s correlation coefficient.

Although non-essential retromer genes were not included in the fitness assays, we first tested our approach on this small and well-characterized complex as a proof of concept, since it has been reported to have PPIs sensitive to gene deletion (Diss et al. 2013). This complex is associated with endosomes and is required for endosome-to-Golgi retrieval of receptors (e.g. the Vps10 protein) that mediate delivery of hydrolases to the vacuole (Mukadam and Seaman 2015). Functionally, the retromer is divided into two subcomplexes: a cargo-selective trimer of Vps35p, Vps29p, and Vps26p and a membrane-bending dimer of Vps5p and Vps17p (Seaman et al. 1998). DHFR-PCA detects significant interactions from all the retromer subunits (*Supplementary Figure S6A*). From our data, we detect same-subunit competition effects between DHFR-fused and non-fused copies of the proteins. Since the extra copy of the duplicated subunit is not tagged with a DHFR reporter fragment, it competes with the tagged copies for the same partners, resulting in a decreased colony size for all the interactions of this subunit (Figure 2A). For instance, we see a reduction of the interaction of all Vps5p PPIs upon duplication of *VPS5* (blue row and blue column, Figure 2A). We also find that the duplication increases the strength of the interactions between Vps35p-Vps26p, members of the other subcomplex, the cargo-selective trimer (Figure 2A). These results show that our strategy has enough resolution to detect small perturbations in the PPIs after the duplication of a single gene coding for a complex subunit.

We next examined the effect of gene duplication on four complexes with proteins for which some gene duplications alter fitness, as identified in our previous analysis, namely the proteasome and the three RNA polymerases (*Supplementary Figure S5*). The proteasome is a highly conserved and thoroughly described eukaryotic protein complex which is amenable to study by DHFR-PCA (Tarassov et al. 2008; Chrétien et al. 2018). In yeast, the core complex (20S) is associated with the regulatory particle (19S) to form a large complex (26S) composed of 37 subunits (Fischer et al. 1994; Hochstrasser et al. 1999). From hereinafter, we will refer to the 26S proteasome as just proteasome. We tested pairwise interactions among 21 subunits as baits and 16 subunits as preys that belong to either the regulatory particle or the core complex for a total of 305 combinations. We detected 47 PPIs between subunits in the *WT* strain (*Supplementary Figure S6B*). The RNA polymerases are also well-described large complexes: RNApol I includes 14 subunits, RNApol II has 16, and RNApol III has 18 subunits (Sentenac 1985; Archambault and Friesen 1993; Cramer 2002). We tested all combinations between all available subunits since five subunits are shared between the three RNA polymerases. We observed 33 significant PPIs out of 689 combinations tested between 31 baits and 26 preys in a *WT* background (*Supplementary Figure S6C*).

We observed same-subunit competition effects (Figure 2B, *Wilcoxon’s rank-sum test p < 2e-16*), which validates that additional copies of the proteins are expressed and that we can measure quantitative changes in their PPIs. Indeed, 135/181 of cases where the duplicated subunit is involved in the PPI tested show a reduced interaction score. We observed that most subunit duplications have small to non-detectable effects on the interaction network of their complex and are weakly correlated with a fitness effect. For the proteasome, excluding competition combinations, only 46 out of 8917 combinations tested were significantly different (FDR of 5%) from the *WT* interactions within the complex. For the three RNA polymerases, only 28 out of 21341 combinations tested were significantly different (Figure 2C). Overall, most of the significant perturbations are gains of PPIs (55/74), which may suggest that when a duplication alters the interaction dynamics within the complex it does so by increasing the strength or amount of PPIs of other subunits (Figure 2C-D). The strongest effects are seen for the duplication of *PRE7*, especially for interactions Pup1p-Pre5p and Pre8p-Rpn8p (Figure 2D). Pup1p, Pre5p, and Pre8p are part of the same subcomplex that includes Pre7p and they interact closely during the formation of the 20S proteasome. Pup1p is the β2 subunit while Pre5p and Pre8p are the α6 and α2 subunits, respectively (Budenholzer et al. 2017) and share close spatial proximity, ranging from 66 to 178Å(Chrétien et al. 2018).

If changes in PPIs are associated with the fitness defects we measure, we hypothesized that we would see a correction between the perturbation of PPIs and fitness effects. We calculated the mean *ps* for each subunit (mean of the absolute values of significant perturbation-scores) and compared it with the selection coefficient of strains containing a gene-duplication of the same subunit (Figure 2E). The correlation between *ps* and fitness is negative as predicted but not significant (*Spearman’s r = −0*.*16, p = 0*.*49*). These results suggest that subunit duplications typically have little or no effect on the protein interaction network within the complex and these effects are largely independent of the fitness effects.

### Most proteasome subunits have an attenuated expression level when duplicated

Our experiments indicate that most PPIs within the proteasome interaction network and RNA polymerases are not significantly perturbed after duplication of their subunits. This suggests that these protein complexes are largely resilient to changes in gene-dosage of their components. Because one of the strongest *ps* was observed in the proteasome, we focused on this complex to explore the underlying mechanisms of such robustness. It has been reported in multiple studies that transcription is usually correlated with gene copy-number while protein abundance correlates more poorly (Dephoure et al. 2014). Therefore, regulatory mechanisms reducing the protein abundance of the proteasome subunits, also known as attenuation, could explain why duplication is not perturbing PPI between the subunits.

To test if the proteasome subunits are attenuated, we looked for changes in protein abundance after gene duplication. We compared protein abundance in GFP-tagged strains (Huh et al. 2003) carrying a duplication of the gene or an empty vector as a control (Figure 3A). Protein attenuation would lead to a reduction of protein levels of both copies, which would result in reduced fluorescence signal. Most subunits (17/19) have a significant decrease in GFP-fluorescent signal after duplication (Figure 3B; *Supplementary Table S6*). Next, we calculated an attenuation score: the difference between *WT* and the duplicated GFP-fluorescent signals normalized by the *WT* (Figure 3C). If the expression of the fused copy was reduced by half to balance the additional copy in a plasmid (complete attenuation), the attenuation score would be 0.5. Interestingly, attenuation is similar for subunits belonging to the same subcomplex (Figure 3D), suggesting a regulation that depends on complex assembly. Havugimana *et* al. (Havugimana et al. 2012) reported that the stoichiometry within each proteasome subcomplex is 1:1, while the stoichiometry among subcomplexes varies from 1:1 to 1:4 (*Supplementary Table S6*). This observation could explain why subunits belonging to the same component have similar expression and attenuation patterns, while there are significant differences between components (Figure 3D; *p = 0*.*002 one-way ANOVA Test*).

**Figure 3.**
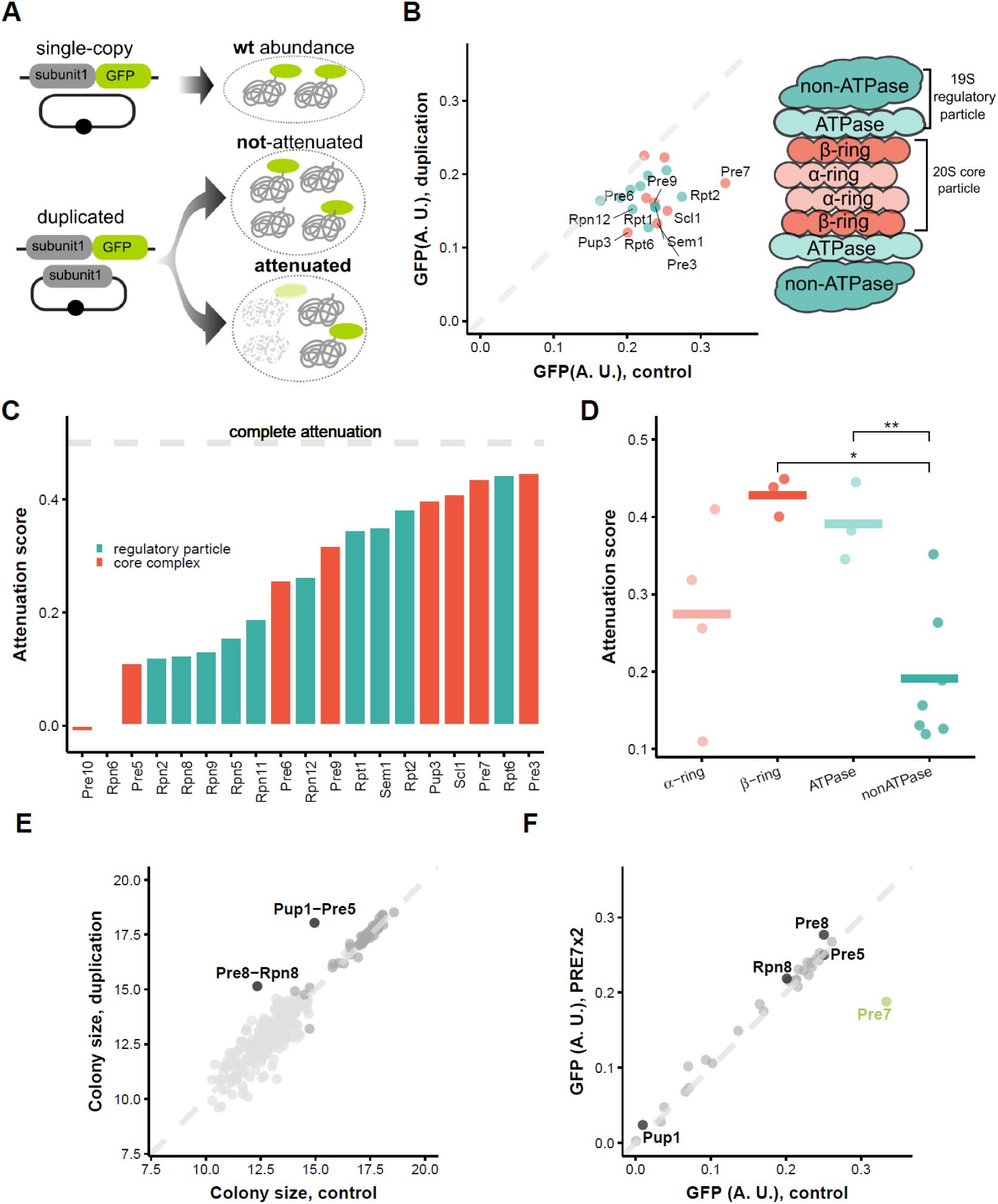
Attenuation of protein abundance after duplication in most proteasome subunits. **(A)** Measuring attenuation with GFP-tagged proteins. Changes in abundance of each subunit can be detected by comparing fluorescent signals of GFP-tagged subunits before and after duplication. Upon attenuation, the abundance of the tagged copy will be reduced. (**B**) GFP signal comparison between strains carrying a duplication of the GFP-tagged subunit and their corresponding control. On the right, a cartoon of the proteasome with its components. All GFP values are corrected for autofluorescence by subtracting the signal of the parental strain not expressing GFP and by cell size (see Methods, *Supplementary TableS6*) (**C**) Attenuation scores of all assayed proteasome subunits. The attenuation score is the difference between GFP fluorescent signals of the control strain (bearing a control plasmid) and the duplicated strain (bearing a centromeric plasmid with an extra copy of the subunit) divided by the GFP signal of the control. In the absence of attenuation, this value is 0. Upon complete attenuation, it is 0.5. (**D**) Attenuation scores of the proteasome subcomplexes. On the right, the asterisks indicate significant differences between components calculated by correcting for multiple testing (*Tukey’s test; * p ≤ 0*.*05 and ** p ≤ 0*.*01*). (**E**) Colony sizes of all strains carrying the *PRE7* duplication (*+pCEN-PRE7*) compared with their control strain (the empty vector) indicating changes in PPI in the DHFR PCA assay. The black dots highlight the subunits that have interactions disturbed after *PRE7* duplication. (**F**) GFP signal of proteasome subunits before and after *PRE7* duplication (*+pCEN-PRE7*). The black dots highlight the subunits that have interactions disturbed after *PRE7* duplication (*Supplementary TableS7*). Replicate measurements are available in the supplementary material.

Strikingly, all but two complex subunits appear to be attenuated. One of the most attenuated subunits Pre7p has one of the strongest effects on both PPIs and fitness when duplicated (Figure 2C). The duplication of *PRE7* perturbs PPIs between Pre5p, Pre8p, Pup1p, and Rpn8p (Figure 3E). All of them share close spatial proximity with Pre7p in the proteasome core complex. We tested if *PRE7* duplication affects the abundance of these subunits by measuring the GFP signal of all proteasome subunits with and without *PRE7* duplication. Most subunits are unaffected upon *PRE7* duplication but the four ‘perturbed’ subunits (Pre5p, Pre8p, Pup1p, and Rpn8p) show a modest increase in their protein abundance (Figure 3F). Even though the difference between the control and the *PRE7* duplicated background is small, it is highly reproducible and significant (*Supplementary Figure S7A*), and appears specific to the subunits with altered interactions (*Supplementary Figure S7B*). These modest but significant changes in protein abundance of ‘perturbed’ subunits after the duplication of *PRE7* may explain the changes we observed in their PPIs. The duplication of *PRE7*, even if largely attenuated, affects the organization of the proteasome by affecting the abundance and interactions of a few other subunits. The dosage balance hypothesis may therefore apply to a limited number of subunits.

The proteasome subunits show different levels of attenuation and recent studies (Dephoure et al. 2014; Chen et al. 2019) suggest that attenuation occurs mostly at the posttranscriptional level across complexes. To examine this specifically for the proteasome, we retrieved data from Dephoure *et al*. (Dephoure et al. 2014) and looked at the mRNA abundance ratio of individual genes in disomic strains relative to wild-type (*WT*). Most proteasome subunits roughly double their transcript levels when duplicated (relative to a log2 mRNA ratio around 1; *Supplementary Figure S8A*). This includes Pre3 and Pre7, two of the most attenuated subunits in our experiments (Figure 3C). Since aneuploidies can cause systemic level changes on the transcriptome and proteome and confound the effects of an individual duplication. We performed RT-qPCR analysis of four highly attenuated subunits (*PRE7, RPT2, RPT6*, and *SCL1*), including three not measured in Dephoure, a non-attenuated subunit (*PRE10*), and a gene that is not a part of the proteasome (*FAS2*). As expected, the non-attenuated genes *PRE10* and *FAS2* present no significant change in their transcript levels after their duplication (*t-test, p = 0*.*6 and 0*.*78, Supplementary Figure S8B*). Highly attenuated genes at the protein level exhibit varying responses: *RPT6* shows a significant mRNA attenuation (*t-test, p = 0*.*01*) and *RPT2* and *SCL1* also show a marginally significant mRNA attenuation *(t-test, p = 0*.*1* and *0*.*07*). *PRE7* displays no significant change in transcript levels (*t-test, p = 0*.*24*). These results from individual gene duplication are therefore consistent with the data observed for disomic strains. Combined, these data suggest that attenuation can occur at the transcriptional level but more frequently at the posttranscriptional level (*Supplementary Figure S8C)*. In most cases, as for *PRE7*, attenuation appears to be posttranslational because transcription and translation rates are both maintained in disomic strains *(Supplementary Figure S8A, (Taggart and Li 2018)*. We further explored the attenuation of the *PRE7* subunit by using a tunable expression system. We constructed a strain with tunable *PRE7* expression (Aranda-Díaz et al. 2017) and which contains two gene copies of *PRE7* tagged with different fluorescent proteins that were independently monitored by flow cytometry. There is a highly significant negative correlation (*Pearson’s correlation r = −0*.*26, p < 10e-15*) between the protein abundance of the copy expressed under the inducible promoter and abundance of the chromosome copy (*Supplementary Figure S9, TableS8*). These results suggest that attenuation is expression level dependent. Finally, we examined whether this posttranslational control depends simply on the presence of an additional gene copy and on that gene copy being transcribed and translated. For this purpose, we constructed a centromeric construction that contains *PRE7*’s full promoter and ORF but that cannot be translated because it lacks a start codon or the start codon is followed by stop codons (*Supplementary Figure S10*). Pre7p remains at the *WT* level of expression in the presence of these three constructions. Since all the *cis*- regulatory elements remain intact on these constructions, it is fair to assume that transcription is initiating but translation is not initiated or is terminated prematurely due to the insertion of the stop codons. This confirms that attenuation requires the translation of the mRNA.

## Discussion

The long-term fate of gene duplicates has been studied in detail theoretically, experimentally and by using genome, transcriptome, and proteome data. While less is known about the immediate impact of gene duplication, it has become clear that dosage sensitivity determines whether gene duplications have any chance to be retained and fixed in a population (Rice and McLysaght 2017a). By duplicating 899 genes individually and examining the distribution of fitness effects, we find that duplicates with a greater than 1% fitness effect are common (∼12%). Furthermore, duplications are twice as likely to be deleterious than beneficial, and deleterious effects are larger in magnitude. Consistent with previous observations, deleterious duplications are more frequent among genes that are also sensitive to a reduction in gene dosage. However, duplications do not have more deleterious effects when they affect protein complexes, contrary to what is predicted from the dosage balance hypothesis. To elucidate this discrepancy, we looked at the effect at the PPI level. To test whether these deleterious effects impact fitness by affecting PPIs in protein complexes, we measured the perturbation of protein complexes *in vivo* as a response to gene duplication, focusing on the proteasome and three RNA polymerases. Overall, only 0.24% of the tested duplication-PPI combinations significantly perturbed their protein complexes, and a single subunit largely drives these results. By focusing on the proteasome, we further examined why so few PPIs were perturbed by changes in gene dosage and found that most of its subunits are attenuated at various extents, i.e. the protein level decreases close to a normal level even if the gene is duplicated or its expression is modified with a tunable promoter. Therefore, our results suggest that gene duplication is unlikely to have an impact on fitness through the perturbation of protein complex assembly at least partly because the extra copies of the genes show attenuated responses at the transcriptional, posttranscriptional and posttranslational levels. Altogether, these observations challenge the dosage balance hypothesis. Our results rather bring support to a model in which decreased dosage, and not increased dosage, affects protein complexes (Semple et al. 2008) by identifying a potential mechanism for this asymmetry of effect. A better understanding of attenuation at the molecular level in the future will allow us to manipulate it in the future and test its causal role in buffering fitness and other molecular effects on the cell.

There is at least one gene that appears to be an exception, as despite being largely attenuated it is highly deleterious and affects PPIs. Further experiments suggest that these alterations are a result of changes in protein abundance of other subunits, which may occur through stabilizing interactions. In previous experiments where we combined gene deletion with the study of PPIs, we documented several cases of protein destabilization by the deletion of an interaction partner (Diss et al. 2013; Diss et al. 2017). What we see here could be the reciprocal effect. Given that Pre7p is one of the most abundant subunits, attenuation may not be sufficient in this case to eliminate the effects of its duplication.

Small-scale duplication (SSD) events of individual ORFs with their cis-regulatory region is less common than other mechanisms of duplication. Indeed, consecutive tandem duplications represent less than 2% of the yeast genome and are not conserved (Despons et al. 2010). For instance, when duplication occurs by retrotransposition, a single CDS is duplicated without its cis-regulatory region (Dujon 2010). Most duplications occur due to recombination errors that lead to the duplication of long segments or even complete chromosomes, leading to the duplication of more than a single gene (Zhang 2003; Dujon 2010). These observations limit the generalization of our observations made on individual duplications. Nevertheless, it has been suggested that the adaptive value of some aneuploidies can be mapped to a single or a handful of loci (Pavelka, Rancati, and Li 2010; Yona et al. 2012). This is the case of the duplication high-affinity sulfate transporter *SUL1* that confers an advantage under sulfate limitation condition (Gresham et al. 2008). Even though our experimental strategy may not fully emulate the most common duplication events in nature, it is a powerful approach to systematically evaluate the impact of individual genes and the possible role of natural selection in the fixation or loss of newly arisen duplicated genes, independently of their origin. In addition, to understand how more complex duplication may impact cell biology and fitness, we first need to understand how individual genes affect cell biology in the first place.

Attenuation of multi-protein complex members has been documented previously in overexpression experiments (Ishikawa et al. 2017), in aneuploid yeast strains (Dephoure et al. 2014; Chen et al. 2019), and to some extent in cancer cells (Gonçalves et al. 2017) which may suffer from a general alteration of protein homeostasis and protein quality control (Oromendia and Amon 2014). Here we show that attenuation is also taking place with small copy-number variation affecting individual genes, such as in the case of gene duplication. This feature appears to be part of the regulation of proteins in normal cells. Indeed, a recent study by Taggart *et al*. (2020) suggested that nearly 20% of proteasome proteins are overproduced since more than half of the protein synthesized is degraded in normal conditions. Therefore, our results suggest that mechanisms acting to regulate protein abundance in normal conditions provide, as a side effect, the extra advantage of protecting the cell against copy-number variation of some members of important multi-protein complexes. In the case of aneuploid cells and protein overexpression, several mechanisms for the attenuation of protein levels have been proposed such as autophagy, the *HSF1*/*HSP90* pathway (Oromendia et al. 2012; Donnelly et al. 2014), and the ubiquitin-proteasome system (Torres et al. 2010; Stingele et al. 2012; Dephoure et al. 2014; Ishikawa et al. 2017). Since the proteasome itself may play an essential role in attenuation, studying the mechanism of attenuation in this complex may prove to be challenging. Nevertheless, the mechanisms protecting the cells of the proteotoxic stress caused by aneuploidy may not be the same that act in the case of SSDs. The molecular mechanisms that attenuate protein abundance in the case of individual gene duplications are still an open question.

Our observations, along with those of previous studies, have an important impact on our understanding of the evolution of protein complexes. Complexes are often composed of multiple pairs of paralogs (Musso et al. 2007; Pereira-Leal et al. 2007), particularly but not exclusively, those coming from a whole-genome duplication (WGD). The relationship between WGD and the retention of gene duplicates in protein complexes has often been explained by a model in which the deleteriousness of single duplications comes from the alteration of protein stoichiometry, which is not affected upon WGD (Papp et al. 2003). However, cells appear to have mechanisms to maintain the stoichiometry of at least some protein complexes, which means that the negative impact of duplication would not come from the perturbation of stoichiometry, but rather from the degradation of excess proteins or other effects not directly related to complex assembly. It is difficult to imagine that such cost, rather than perturbations of stoichiometry, would explain the observation that complex subunits are more likely to be retained after a WGD event, but are less likely to be duplicated individually (Papp et al. 2003; Qian and Zhang 2008). This explanation, therefore, needs further examination and the extension of the study of attenuation to a larger number of complexes.

Evolutionary forces leading to regulatory mechanisms that attenuate protein abundance, therefore, could diminish the immediate fitness effects of gene duplications and could allow them to reach a higher frequency in populations. Such pressures to maintain gene dosage could come, for instance, from a constant requirement to assemble protein complexes in a stoichiometric fashion in the face of gene expression noise (Fraser et al. 2004). Another pressure for the degradation of extra-subunits is the need to prevent spurious interactions between the unassembled subunits and other proteins through its exposed sticky interface (Levy et al. 2012) or aggregation (Brennan et al. 2019). If selection for decreasing noise or to reduce spurious interactions led to the evolution of expression attenuation, it may have also contributed to the robustness of protein complexes to gene duplications. Then, if the cost of producing extra RNA and protein is not too high, newly arisen duplicated subunits would have a higher probability of fixation and of long-term maintenance, perhaps by dosage subfunctionalization (Qian et al. 2010; Gout and Lynch 2015).

## Material and methods

### Parental strains, media, and plasmids for the fitness measurements

We extracted the MoBY plasmids from bacteria using a 96-well plate miniprep extraction by alkaline lysis (Engebrecht et al. 1991). The parental strain used for the fitness assay library is Y8205-mCherry (*MATα ho::PDC1pr-mCherry-hphMX4 can1Δ::STE2pr-Sp_his5 lyp1Δ::STE3pr- LEU2 his3Δ1 leu2Δ0 ura3Δ0*), a modification of the Y8205 strain from (Tong and Boone 2006) done by (Garay et al. 2014). The reference strain used for the competition assays was Y7092- CFP (*MATα hoΔ::Cerulean-natR can1Δ::STE2pr-SP_his5 lyp1Δ Δhis3Δ1 leu2Δ0 ura3Δ0 met15Δ0 LYS2+*) transformed with a control plasmid (p5586; (Ho et al. 2009)) identical to those from the MoBY-ORF collection but without yeast gene sequence. To generate the collection with duplication of essential genes, we used a high-efficiency yeast transformation method for 96-well format (Burke et al. 2000). To select positive transformants we incubated plates at 30°C for 3-5 days and cherry-picked two colonies per sample. The competition medium was a minimal low-fluorescence synthetic medium without uracil (SC[MSG]-lf -ura, *Supplementary TableS10*).

### Measurements of relative fitness by fluorescence-based competition assays

To measure relative fitness we used an automated high-resolution method based on fluorometry previously developed by (DeLuna et al. 2008). The mCherry-tagged collection carrying individual ‘duplications’ (Y8205-mCherry +MoBY-xxx) were competed with a universal CFP-tagged strain (Y7092-CFP +p5586). First, we inoculated 150 µL of fresh medium (using Corning® Costar® 96- well cell culture plates) with overnight cultures of ‘duplicated’ and reference strains (tagged with mCherry and CFP respectively) in a 1:1 proportion. Every 24 h, we diluted the cultures 16-fold into sterile fresh competition media. Cultures were monitored in parallel for approximately 28 generations (7 days) in a fully automated robotic system (Freedom EVO, Tecan Ltd) connected with a microplate multi-reader (Infinite Reader M100, Tecan Ltd). We monitored OD600 nm, mCherry fluorescence signal (578/610 nm, gain: 145), and CFP fluorescence signal (433/475 nm, gain: 120) every 2 h. Cultures were incubated at 30 °C and 70 % relative humidity.

### Estimation of selection coefficients

We estimated the selection coefficient by measuring the change in population ratios of cells carrying a duplication (mCherry signal) on the reference strain (CFP signal), following the standard procedures developed by (DeLuna et al. 2008). Each fluorescent signal was corrected for media autofluorescence and for the crosstalk between fluorescent signals. To calculate each fluorophore crosstalk, we measured mCherry_BKG_ background signal in CFP-only wells and CFP_BKG_ in mCherry-only wells. Then, we subtracted the background signals: mCherry = mCherry_RAW_- mCherry_BKG_ and CFP = CFP_RAW_-CFP_BKG_. Corrected fluorescence values were used to estimate *ln(mCherry/CFP)* ratios for all measurement points. Then, we interpolated a single ratio per dilution cycle at a fixed OD600nm (0.2) to reduce artifacts related to the dependence between the fluorescent signals. We discarded any extreme ratios (*ln(mCherry/CFP)* > 4 or < −4) and samples with less than three data points. We calculated the selection coefficients using the equation:

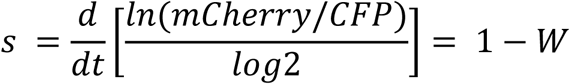

Where: ***s*** is the selection coefficient, ***t*** is the number of generations, and ***W*** is relative fitness. We normalized each *s* using the median of the complete distribution of strains. We calculated z-score using the mean and the standard deviation of a control distribution of 192 Cherry-tagged *WT* strains competed against the universal CFP-tagged reference. All calculations were performed using scripts written in MatLab and R.

### Measurements of relative fitness by cytometry-based competition assays

The Y8205-mCherry +MoBY-xxx collection was co-cultured with a universal YFP-tagged strain (Y7092-YFP +p5586). Daily dilutions steps were carried as the fluorescence-based competition assays. Before each dilution, we took a sample of 10 μL of saturated culture and made a 1:10 dilution in TE 2X (*Supplementary Table S10*) in 96-well plates. Cytometry measurements were performed on a LSRFortessa™(BD, USA) cytometer. We collected up to 30,000 events using the integrated high-throughput sampler. We calculated the number of cells expressing mCherry or YFP fluorescence using the FACSDiva™ (BD, USA) software tool for manual gating. Selection coefficient estimation was done as described above.

### Measurements of relative fitness in different conditions

We confirmed by PCR the MoBY plasmid identities and transformed them into *PDC1*-mCherry strain. From each transformation, we isolated four independent colonies as replicates to measure relative fitness. The reference strain (*PDC1*-mCherry+pRS316-KanMX) was competed with another reference carrying a control empty vector (*PDC1*-GFP +pRS316-KanMX) to have a reference versus reference control. We grew individual pre-cultures of all strains for 48 hours at 30 °C in the standard media (SC -ura). We then mixed 120 uL of saturated culture of the *PDC1*- mCherry +MoBY strains with 100 uL of the reference *PDC1*-GFP +pRS316-KanMX. We used 20 uL from these mixtures to inoculate 500 uL of fresh media. We tested five different media: SC - ura, SC -ura +1.2 M Sorbitol, SC -ura + 2mg/mL Caffeine, SC -ura + 6% Ethanol and SC -ura with 2% Galactose as a carbon source. Plates were sealed with breathable sterile seals and incubated at 30°C at 400rpm. Every 24 h we diluted the saturated cultures 1:20 into fresh media of the same composition. Every day, before dilution, we took a sample of the saturated culture and adjusted to 500 cells/uL in water to measure relative cell count in the cytometer. We followed the cultures through 6 dilution cycles.

### Construction of MoBY-EGFP plasmids by Gibson Assembly

We generated pCEN constructs coding for protein fusions with EGFP using Gibson Assembly (Gibson et al. 2009). First, we amplified the EGFP tag (insert fragment) from the plasmid pFA6- GFP(S65T)-HIS3MX using oligonucleotides targeting this ORF without selection markers. In parallel, we amplified the pCEN backbone with oligonucleotides targeting the ORF without stop codon (*Supplementary Table S9*). Both PCR reactions were digested with DpnI (NEB, USA) for one hour at 37°C and then purified using the MagJET NGS Cleanup Kit (Thermo Fisher Scientific, USA). 50 ng of the insert and 50 ng of the backbone (molar ratio of ∼1:10) were adjusted to 2.5 µL and mixed with 7.5 µL of Gibson assembly master mix prepared accordingly with manufacturer instructions (NEB, USA). Reactions were incubated for 1 hour at 50°C. We transformed BW23474 (F−, Δ*(argF-lac)169*, Δ*uidA4::pir-116, recA1, rpoS396 (Am), endA9(del-ins)::FRT, rph-1, hsdR514, rob-1, creC510*) competent cells with 5µL of each reaction and selected on 2YT plates supplemented with 50 mg/L Kanamycin. We confirmed 8 colonies per transformation by colony PCR and subsequent Sanger sequencing of the purified plasmids. These confirmed plasmids were transformed into BY4741 competent yeast cells and selected on SC -ura media. Six independent colonies from each transformation were isolated for further analysis.

### Measuring protein abundance of Act1 by Western Blot assay

Strain PGK1-GFP from the Yeast GFP fusion Collection (Huh et al. 2003) was transformed with pRS316-KANMX or MoBY-*ACT1* plasmid. Preculture of 3 independent colonies of each transformation was performed in 5mL of SD[MSG] -ura -his media for 16h at 30°C with agitation. The next day, precultures were diluted at 0.05 OD/mL in 100mL of fresh SD[MSG] -ura -his media, let grow to ∼0.5 OD/ml and at the end 20U OD was spun and frozen at −80°C. Cell pellets were resuspended in 200 µl of water with peptidase inhibitors (Sigma Aldrich, USA). Glass beads were added and the suspension was vortexed for 5 min. SDS 10% was added (final concentration 1%) and the samples were boiled for 10min. Samples were clarified by centrifugation at 13,200 rpm for 5min. The equivalent of 0.5U of OD was separated on a 10% SDS-PAGE and transferred to a nitrocellulose membrane. *PGK1-GFP* was detected with an anti-GFP (Sigma Aldrich, USA) and actin was detected with an anti-actin clone C4 (Millipore, USA). Primary antibodies were detected with a goat anti-mouse 800 (LI-COR Biosciences, USA). The membrane was imaged using an Odyssey Fc Imaging System (LI-COR Biosciences, USA). PGK1-GFP was used as the loading control and actin amount was compared between strains expressing pRS316-KANMX and those expressing MoBY-*ACT1*.

### Generation of DHFR-PCA strain library with duplicated subunits

We retrieved strains that contain DHFR fragment fusions of subunits of the proteasome, RNA polymerase I, II, and II from the Yeast Protein Interactome Collection (Tarassov et al. 2008). We confirmed the correct insertion of the DHFR fragments by colony PCR using an ORF specific forward Oligo-C (Giaever et al. 2002) located upstream of the fusion and a universal reverse complement oligonucleotide ADHterm_R (*Supplementary Table S9*). PCR amplicons were sequenced to confirm correct fusion with the DHFR fragments. All oligonucleotide sequences can be found in *Supplementary Table S9*. PCR reaction conditions and reagents are the same as in (Marchant et al. 2019). The number of confirmed DHFR strains is indicated in *Supplementary Table S4*.

The 70 plasmids that were retrieved from the MoBY-ORF collection (Ho et al. 2009) contain subunits from the Retromer (6), Proteasome (33), and RNA polymerases I, II and II (31) (see *Supplementary Table S4)*. We confirmed their identities using Oligo-C and a universal plasmid-specific oligo KanR (*Supplementary Table S9*). All plasmids were extracted using the Presto Mini Plasmid Kit (Geneaid Biotech Ltd, Taiwan).

The strains expressing DHFR fusions and an additional gene copy from the MoBY plasmids or the empty plasmid were transformed following a standard protocol (Burke et al. 2000). The selection was done in three steps. The first round was done in 96 deep-well plates in 800 μL of SD[MSG] -ura, incubated for 2 d at 30°C with shaking. The second round was done by inoculating 5 μL of the first-round selection cultures into 800 μL of fresh selection media. Finally, cells were deposited on agar plates with double selection SD[MSG] -ura + G418 (200 mg/L).

We took all confirmed preys (DHFR-F[3] fusions) and transformed them with all MoBY plasmids available for their complex (*Supplementary TableS4*). We generated 42 strains with the 6 retromer preys each transformed with seven plasmid constructions (6 MoBY-xxx plasmids plus one control plasmid). For the proteasome, 544 strains were constructed by transforming each of the 16 preys with the 34 plasmids (33 MoBY-xxx plasmids and one control plasmid). For the RNA polymerases I, II, and II, we generated 864 strains, 27 preys, and transformed them with 32 plasmids (31 from the MoBY collection and one control plasmid). The control plasmid (pRS316-KanMX) contains the same selection markers as the MoBY plasmids to account for the effect of expressing the markers alone.

### DHFR Protein fragment Complementation Assay

DHFR-PCA detects semi-quantitative changes in strength and frequency of the physical interaction between two proteins (Freschi et al. 2013; Levy et al. 2014). This method allows the detection of PPI changes upon perturbations *in vivo*, for instance in response to gene deletion (Tarassov et al. 2008; Diss et al. 2013; Diss et al. 2017). To measure the effect of gene duplication, we introduced an extra copy coded on a centromeric plasmid to simulate gene duplication as above, and measured changes in the physical interaction between pairs of the complex subunits. The DHFR-PCA screening was performed using standard methods used in previous works (Rochette et al. 2015; Diss et al. 2017; Marchant et al. 2019). All arrays of 96, 384, and 1,536 colonies per plate were manipulated with pin tools controlled by a fully automated platform (BM5-SC1, S&P Robotics Inc., Canada; (Rochette et al. 2015)). Haploid strains were assembled in arrays of 1,536 colonies on agar plates. In order to avoid growth bias at the edge of the agar plates, the border positions (first two and last two columns and rows of each 1,536 array) of all plates were dedicated to positive controls (*MAT***a** *LSM8*-DHFR-F[1,2] and MAT**α** *CDC39*-DHFR-F[3]). These two proteins have previously shown a strong interaction in PCA (Marchant et al. 2019).

The complete *MAT***α** *prey-DHFR-F[3] +MoBY-xxx* (prey) collection of 1,450 strains was expanded to generate seven independent replicates using a cherry-picking 96-pin tool. Each replica was positioned randomly in the 384-colony array, avoiding border positions. During the cherry-picking step of the preys, we generated 34 plates with 384-colony arrays. We condensed them into 9 plates (one for the retromer, three for the proteasome, and five for the RNA polymerases) of 1,536-colony arrays. All prey manipulations were performed on YPD +hygromycin B (250 mg/L) and G418 (200 mg/mL) media.

For the *MAT***a** *bait*-DHFR-F[1,2] (bait) collection, we produced 52 arrays of 1,536 colonies printed on YPD +NAT (100 mg/L) agar plates. On the interior positions, each plate contained the same bait-strain of the Proteasome (21) and the RNA polymerases (31), while in the border positions, we printed the same control strain. In the case of the retromer, six 384-colony arrays were condensed in two plates with 1,536 colonies. As a result, we produced 54 PCA-ready bait plates.

The 9-plate prey collection (*MAT***α** *prey*-DHFR-F[3] +MoBY-xxx, Hygromycin B, and G418 resistant) was mated with the set of 54 bait plates (*MAT***a** bait-DHFR-F[1,2], NAT resistant). All crosses were performed within each complex in the following manner: for the retromer prey plate was crossed with two bait plates, generating two mating plates. For the proteasome, three prey plates were crossed with 21 bait plates, generating 63 mating plates. Finally, for the RNA polymerases, each of the five prey plates was crossed with the 31 corresponding baits, resulting in 155 mating plates. Mating was done on agar plates of YPD and incubated at room temperature for 2 d. Then, we replicated the set of 220 mating plates on diploid selection media containing the three antibiotics (YPD +hygromycin B (250 mg/L), NAT (100 mg/L), and G418 (200 mg/L) and incubated the plates at 30 °C for 48 h. We repeated this step to improve colony homogeneity. Finally, the complete set of diploids was replicated on both DMSO and MTX media (*Supplementary Table S10*) two consecutive times, each step with incubation at 30 °C for 96 h. We monitored growth in the last step of both DMSO and MTX selections. We captured images of the plates every 24 h using a Rebel T5i camera (Canon, Tokyo, Japan) attached to our robotic system (S&P Robotics Inc, Toronto, Canada).

### Data Analysis of the DHFR-PCA screening

We used a plate image analysis method to measure colony sizes as the integrated pixel density of each position of the 1,536-array plates. Plate pictures were processed by a customized pipeline that was implemented using the ImageJ 1.45 (http://rsbweb.nih.gov/ij/) software as described previously by (Diss et al. 2013; Diss et al. 2017). We calculated a PPI-score for each position by subtracting colony size on the control medium (DMSO) to the colony size on the DHFR selective media (MTX) to eliminate any fitness or position bias. Then, we calculated a mean PPI score by averaging the specific interaction scores of the seven biological replicas. We used the R function ‘normalmixEM()’ (Benaglia et al. 2009) to adjust the PPI scores to a bimodal distribution. This model was used to determine a PPI score threshold of 14.7 for positive interactions. We excluded all positions below this threshold and samples with less than five biological replicates, leaving 90% of samples after filtering. We calculated *ps* as the difference of PPI scores between ‘duplicated’ (+MoBY-xxx) and *WT* (+pRS316-KanMX) background. We estimated statistical significance with a two-sided paired Student’s T-test. The resulting p-values were then adjusted (to q-values) for multiple comparisons using the FDR method (Benjamini and Yekutieli 2001) with the p.adjust() function in R. Samples with q-values below 0.05 were considered positive hits. All the analysis pipeline was done using custom R scripts.

### Measurements of protein abundance by flow cytometry

We used GFP(S65T)-HIS3MX–tagged strains from the Yeast GFP fusion Collection (Huh et al. 2003). The identities of the 33 strains expressing GFP-tagged proteasome subunits were confirmed by colony PCR using an ORF specific oligo (*Supplementary Table S9*). We generated *de novo* GFP-fusions for four subunits (*PRE7, RPT2, PRE1*, and *PUP3*) that were not present in the Yeast GFP Fusion Collection. These fusions were produced following the same strategy described in the original work (Huh et al. 2003). We confirmed at least one clone of each new GFP-tagged construction by cytometry, colony PCR, and sequencing (*Supplementary Table S9*). Each GFP-tagged strain was transformed with a plasmid of the MoBY ORF library expressing an additional gene-copy of the same tagged subunit. Since there are only 33 subunits present on the MoBY ORF library, therefore, attenuation could only be measured for the corresponding proteins.

For the cytometry measurements, we inoculated the strains in a 96 deep-well plate in 500 µL of SD[MSG] -ura -his media. The plate was sealed with a sterile breathable membrane. After overnight growth at 30 °C (with shaking), the saturated culture was diluted 1:10 in the same media. Cells were incubated again for 4-6 hours at 30°C with agitation. Then, cells in exponential phase were diluted in sterile ddH_2_O to an approximate concentration of 500 cells/µL. The GFP fluorescence of 5000 cells was measured using a Guava easyCyte 14HT BGV flow Cytometer (Millipore, USA). All data processing was done using custom R scripts. Events with FSC-H or SSC-H values below 1000 were discarded. The GFP signal for each event was normalized with cell size using the FSC-A value. Finally, we subtracted the autofluorescence value of the cells, calculated by measuring the parental BY4741 strain without any fluorescent protein. For further analysis, we only considered strains with GFP signals above the median plus 2.5 standard deviations of the *WT* non-GFP strain.

### mRNA quantification by RT-PCR

mRNA abundance was measured using quantitative real-time PCR (qRT-PCR) in strains expressing (MoBY plasmid) or not (pRS316) an additional copy of the gene of interest. Total RNA was extracted from cells grown to OD600 0.4–0.6 in 6 mL of SD[MSG] -ura -his using standard hot-phenol procedure (Köhrer and Domdey 1991) and RNA samples were treated with DNase I (New England BioLabs, USA) according to manufacturer’s protocol. RNA yield was measured using a Nanodrop 2000c spectrophotometer (Thermo Scientific, USA). One μg of total RNA template was used for each cDNA synthesis in reactions including 5 μM oligo d(T)12–18, 1 mM dNTPs and 200 U M-MuLV (New England BioLabs, USA) which were treated following manufacturer’s instructions with some modifications: 1) no RNAse inhibitor was used, 2) reactions were incubated for 10 min at 25 °C prior to cDNA synthesis, 3) reactions were incubated for 50 min at 42 °C for cDNA synthesis and 4) the enzyme was inactivated for 15 min at 70 °C. cDNA products were diluted in three volumes of water and 2 μl was used for qRT-PCR analysis in 2× Real-Time PCR premix or in 2× PerfeCTa SYBR Green FastMix (Quantabio) on a 7500 Real-Time PCR System (Applied Biosystems, USA). A universal PCR primer pair for all the tested genes and specific to the GFP fusion was used (*Supplementary Table S9*). *ACT1* and *ALG9* were used as control genes. Serial dilutions of plasmids containing the GFP encoding gene (GFP- *ATG8* (Reggiori et al. 2004) *and ACT1* and *ALG9* genes (MoBY-*ACT1* and MoBY-*ALG9*) were used to generate standard curves for the quantification. Up to three qRT-PCR technical replicates from three biological replicates (three different cell cultures, RNA extractions and cDNA synthesis) were performed for each strain. The absence of contaminant genomic DNA was confirmed using a template-negative control and no RT-controls. The quantity of GFP, *ACT1* and *ALG9* mRNA was estimated using the standard curves derived from the reference plasmids and the expression level of each gene fused to GFP was measured as the ratio of GFP to *ACT1* or to *ALG9* abundance.

### Attenuation experiments with an estradiol inducible promoter system

To construct the inducible promoter plasmid, we cloned *PRE7* in the pAG416GAL-ccdB-EGFP plasmid (Addgene plasmid # 14195). To generate this construction, we amplified *PRE7* (ORF only) from the strain BY4741 genomic DNA using oligonucleotides listed in *Supplementary Table S9* that contain gateway attR recombination sequences. We used 300 ng of purified PCR product to set a BPII recombination reaction (5 μL) into the Gateway Entry Vector pDONR201 (150 ng) using the standard methods provided by the manufacturer (Invitrogen, USA). BPII reaction mix was incubated at 25 °C overnight. The reaction was inactivated with proteinase K. The 5 μL reaction was used to transform MC1061 competent *E. coli* cells. The selection was performed in 2YT + 50 mg/L of kanamycin (BioShop Inc, Canada) at 37 °C. Positive clones were detected by PCR using an ORF specific oligonucleotide and a general pDONR201 primer (*Supplementary Table S9*). We isolated plasmid from one positive clone that was confirmed both by PCR and by Sanger sequencing. The LRII reaction was performed mixing 150 ng of pDONR201-PRE7 and 150 ng of pAG416GAL-ccdB-EGFP. The reaction was incubated overnight at 25°C and inactivated with proteinase K. The reaction was used to transform MC1061 competent *E. coli* cells, followed by selection on solid 2YT + 100 mg/L ampicillin (BioShop Inc, Canada) at 37°C. Positive clones were confirmed by PCR and Sanger sequencing using *PRE7* and pG416GAL- ccdB-EGFP specific oligonucleotides (*Supplementary TableS9*). The sequence-verified pAG416GAL-PRE7-EGFP plasmid was used to transform the yeast strain *PRE7*-mKate (BY4741 *MAT*a *Δhis3 Δura3 Δmet15 LEU2::GEM PRE7-mKate-hph*) using a lithium acetate standard protocol. As a reference, we transformed the same parental strain with the empty expression plasmid pG416GAL-ccdB-EGFP. The selection of positive transformants was done on SC[MSG] -ura -leu for 3 d at 30 °C.

For the induction experiments, we set up overnight cultures of the PRE7-mKate + pAG416GAL- PRE7-EGFP strain and the reference (*PRE7*-mKate + pG416GAL-ccdB-EGFP) in 5 mL of SC[MSG] -ura -leu. We diluted the saturated culture 1:10 in 96-well plates containing 200 μL of SC[MSG] -ura -leu containing Estradiol (β-estradiol, Sigma-Aldrich) at different concentrations (0, 6.25, 12.5, 25, 50 and 100 nM). We incubated at 30°C with shaking for 12 hours. Then, cells in exponential phase were diluted in sterile ddH_2_O to an approximate concentration of 500 cells/µL. mKate and GFP fluorescence signals were measured as described previously.

### Mutagenesis of MoBY-*PRE7* plasmid

To generate MoBY-*PRE7* plasmids without the translation of Pre7p, we performed site-directed mutagenesis. For a 25 μL mutagenesis reaction, the following were mixed: 5 μL Kapa HiFi buffer 5X (Kapa Biosystems, USA), 0.75 μL dNTPs 10 μM, 0.75 μL forward oligonucleotide 10 μM, 0.75 μL reverse oligonucleotide 10 μM (see *Supplementary Table S9*), 0.5 μL Kapa hot-start polymerase (Kapa Biosystems, USA), 0.75 μL MoBY-PRE7 plasmid DNA (15 ng/μL), 16.5 μL PCR grade water. The following thermocycler protocol was then used: 95 °C for 5 min, then 20 cycles of 98 °C for 20 s, 60 °C for 15 s, 72 °C for 11 min and then 72 °C for 15 min. The PCR product was digested with DpnI for 2 h at 37 °C and 5 μL was transformed in *E. coli* strain BW23474. Transformation reactions were spread on agar plates of 2YT media with Kanamycin (50 mg/L) and Chloramphenicol (12.5 mg/L) and incubated at 37 °C overnight. Three independent colonies were isolated from each transformation and confirmed by sequencing.

### Databases used for data analysis

The list of genes that code for members of multi-protein complexes of *S. cerevisiae* (S288c) was obtained from the Complex Portal (https://www.ebi.ac.uk/complexportal/home) developed and maintained by (Meldal et al. 2019).

Haploinsufficient genes were taken from the work of (Deutschbauer et al. 2005).

All MatLab and R scripts used for data analysis and visualization are available in a GitHub repository (url: https://github.com/Landrylab/AscencioETAL_2020).

## Acknowledgments

We thank E. Mancera and members of the Landry and DeLuna labs for discussions and comments on the manuscript. This research was supported by a CIHR Foundation grant to CRL (387697) and by CONACYT México through grant PN-2016/2370 to AD. CRL holds the Canada Research in Cellular Synthetic and Systems Biology.

## Abbreviations

DHFR: Dihydrofolate Reductase
GFP: Green Fluorescent Protein
FDR: False Discovery Rate
PCA: Protein Complementation Assay
pCEN: Centromeric Plasmid
PPIs: Protein-protein Interactions
*ps*: Perturbation Score
*s*: Selection Coefficient
WGD: Whole-genome Duplication
*WT*: Wild type

## Supplementary Information

### Supplementary Figures

**Figure S1.**
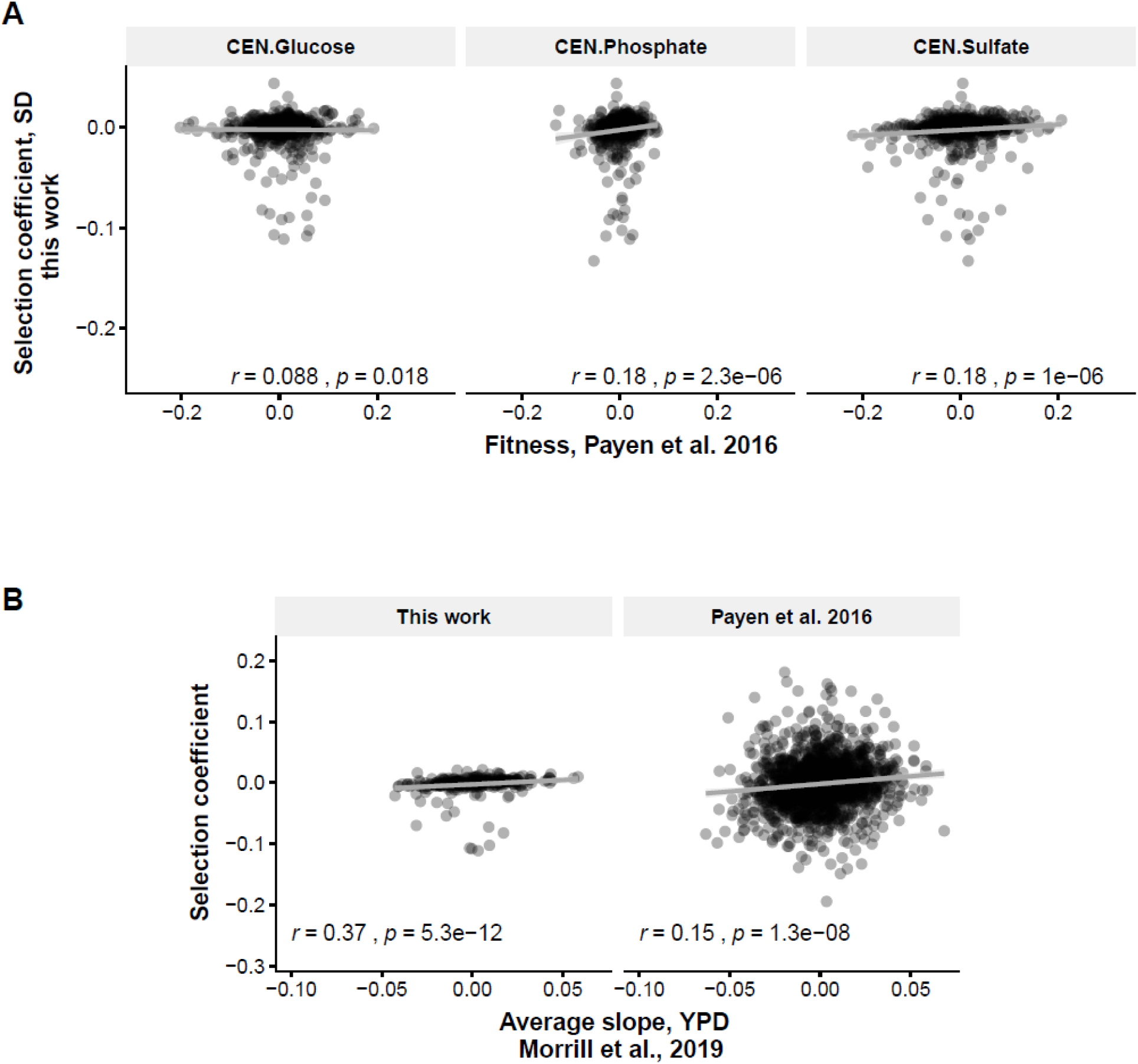
Comparison of the relative fitness measurements with previous studies. (**A**) Comparison of the selection coefficient measured by fluorometry in this work and those of pool assay competitions reported Payen *et al*. (2016) in three different nutrient-limiting conditions (**B**) same comparisons with the pool assay competition in rich media reported by Morril *et al*. (2019). We show Spearman’s correlation and p-value at the bottom.

**Figure S2.**
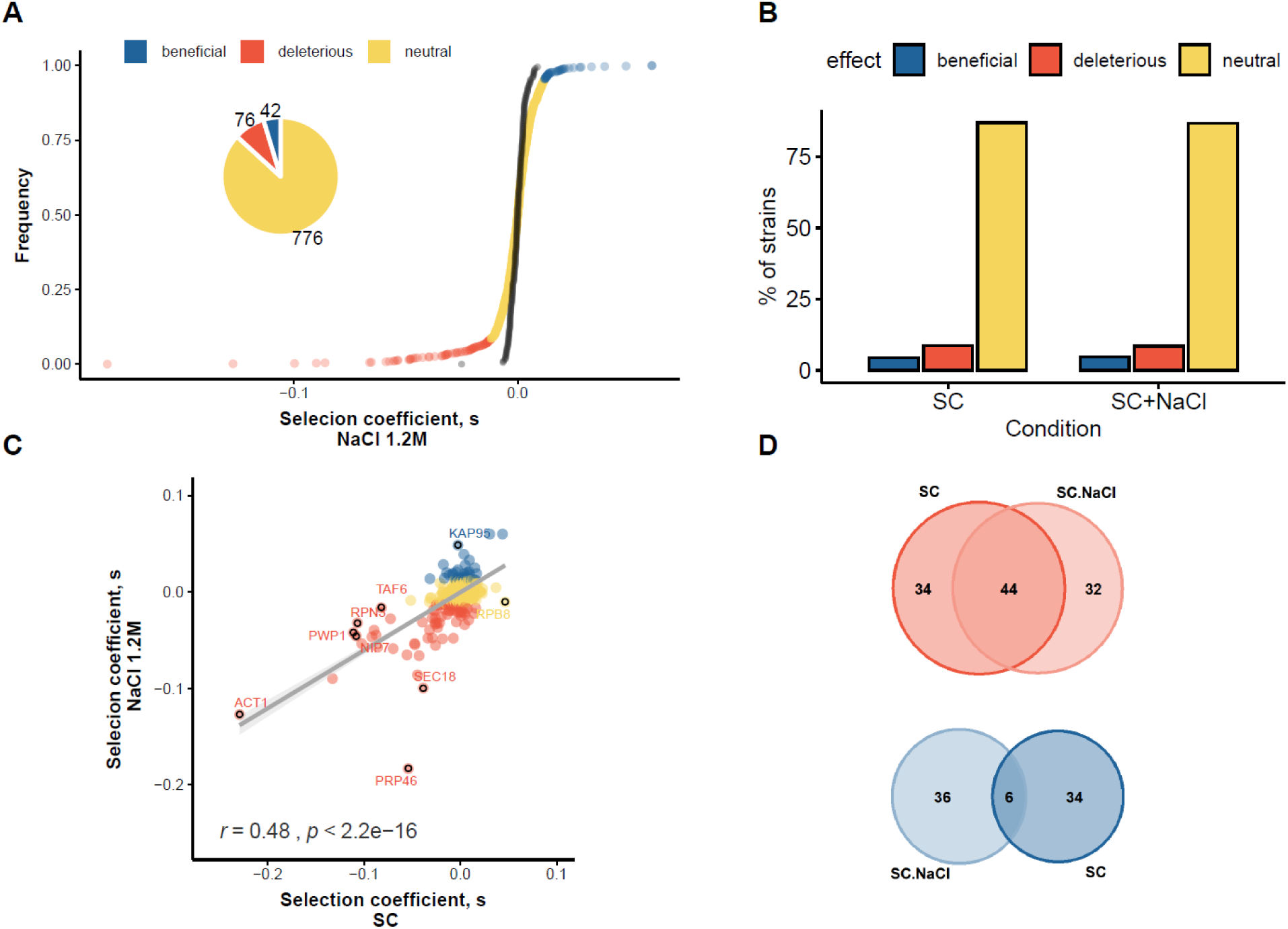
Results from the competition experiment in ionic and osmotic stress. (**A**) Cumulative distribution of the selection coefficients of strains with additional copies of essential genes tested in stress. Experimental conditions are the same as the nominal media. Genes are classified in neutral, deleterious and beneficial using (selection coefficient > 0.01 or < −0.01, |z-score| > 4.5). (**B**) Percentage of strains that are neutral (yellow bars), deleterious (red bars), and beneficial (blue bars) effects in both conditions tested. (**C**) Comparison of selection coefficients in both conditions. Spearman’s correlation coefficient and p-value are shown at the bottom. Black circles highlight strains within more than a 5% difference in relative fitness between the two conditions. (**D**) Venn diagrams for genes with fitness effects in both conditions.

**Figure S3.**
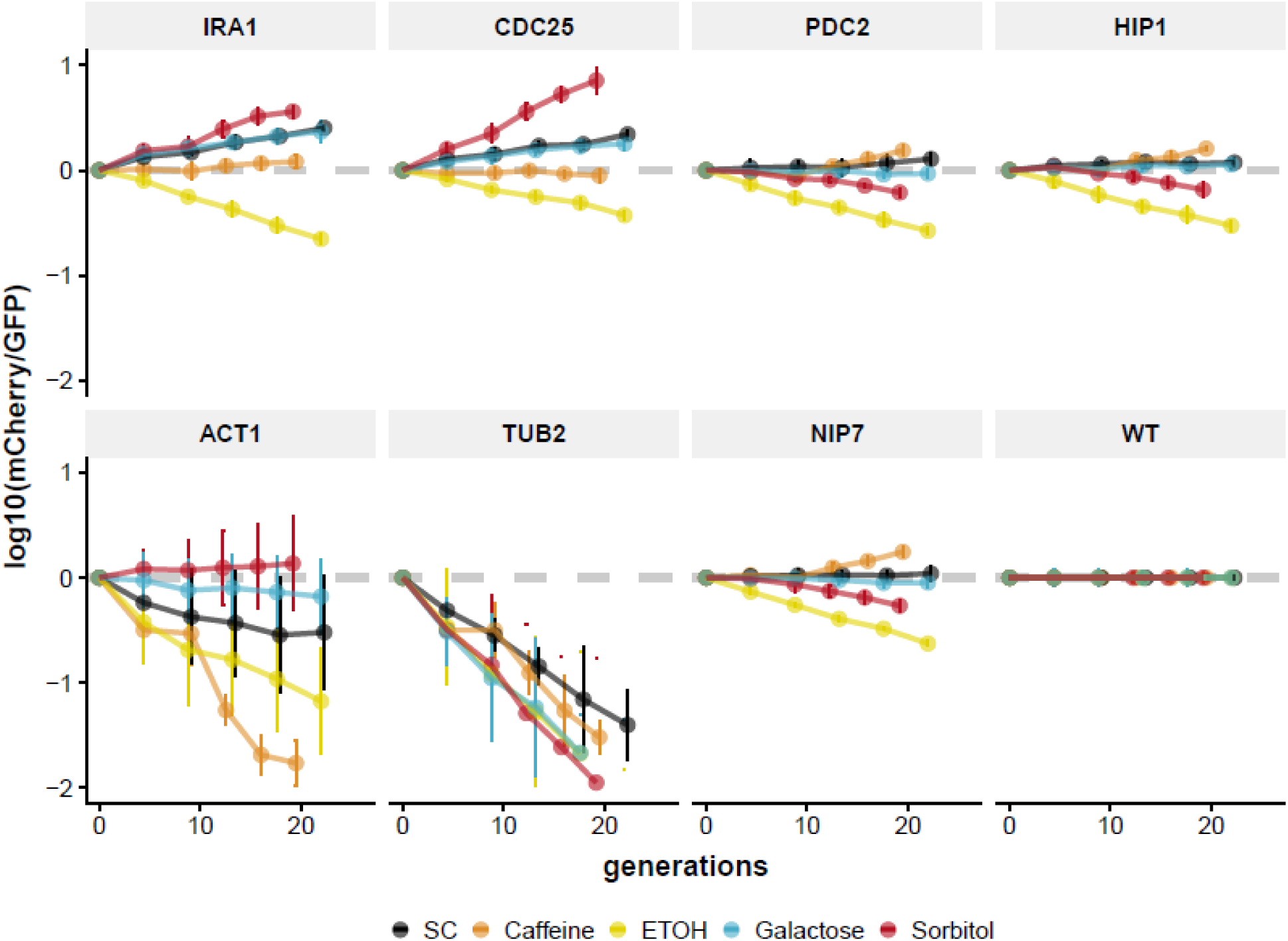
Competitive fitness assay in different growth conditions. Each dot represents the ratio of the number of mCherry cells (+pCEN-xxx) relative to the number of reference GFP cells (+pRS316-KanMX), as a function of the number of generations. Bars represent the standard error of the mean of four biological replicates. The title of each subplot indicates the +pCEN-xxx carried by the mCherry strain. Different colors indicate the condition used in each curve, see methods.

**Figure S4.**
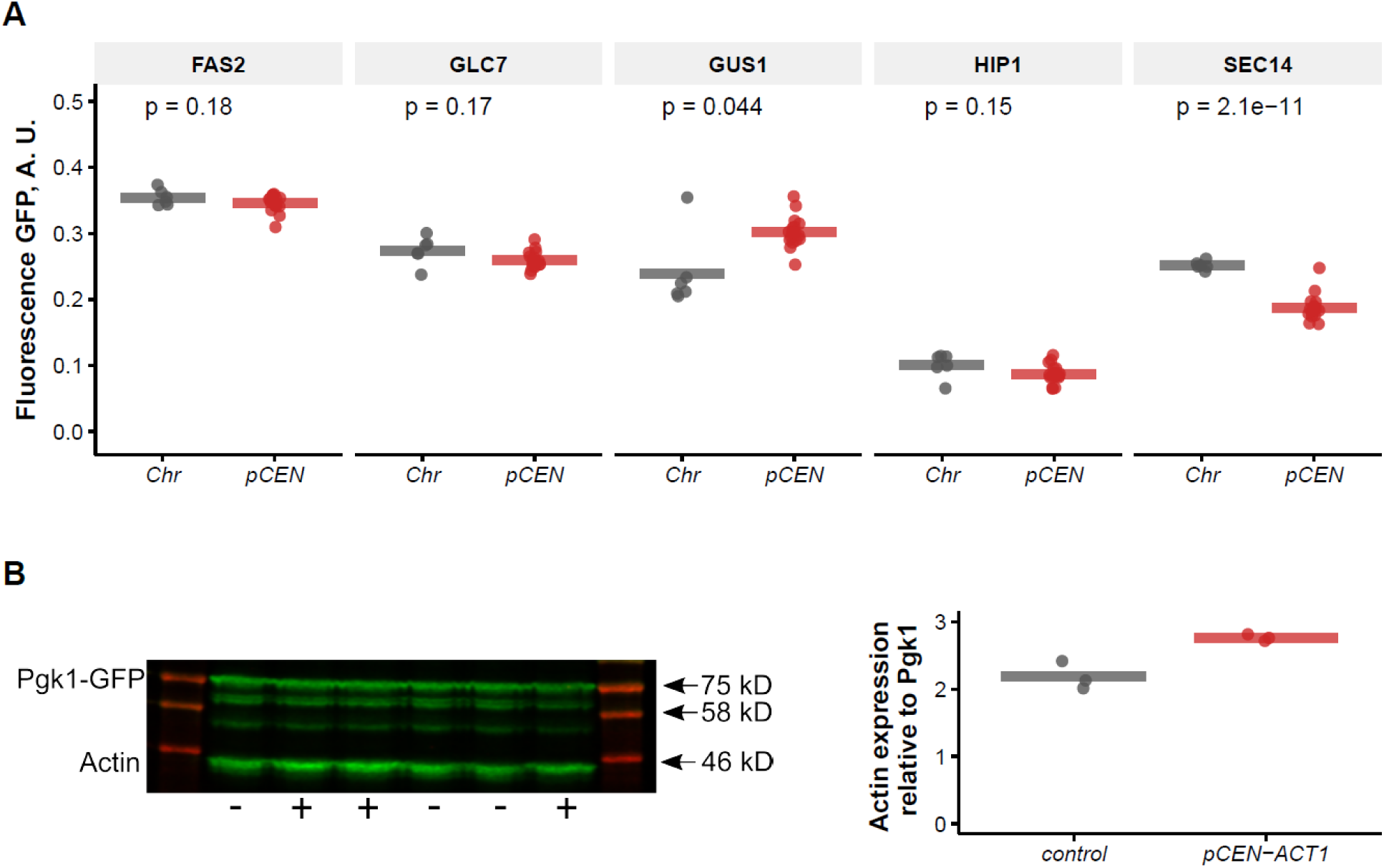
Expression from single-copy plasmids approximates expression from the genome. (**A**) The fluorescent ratios of strains bearing the indicated gene fused with EGFP on its native locus on the chromosome (Chr, grey) and on a pCEN construct (red). In order to discard the effects of adding an extra gene copy, regardless of where the EGFP fusion is coded, all the strains carry two gene copies one in a chromosome and the other in a pCEN. Each dot represents the median of 5000 individual events measured by flow cytometry. (**B**) The protein abundance of Act1p. Western blot analysis showing protein levels of a *WT* strain (-, carrying an empty vector) and a strain carrying pCEN-*ACT1* (+). We show the results for three replicated cultures. Both strains express a Pgk1-GFP. Western immunoblots probed with antibodies against GFP and actin. The quantification of the actin signal over the Pgk1-GFP signal is shown on the boxplot on the right.

**Figure S5.**
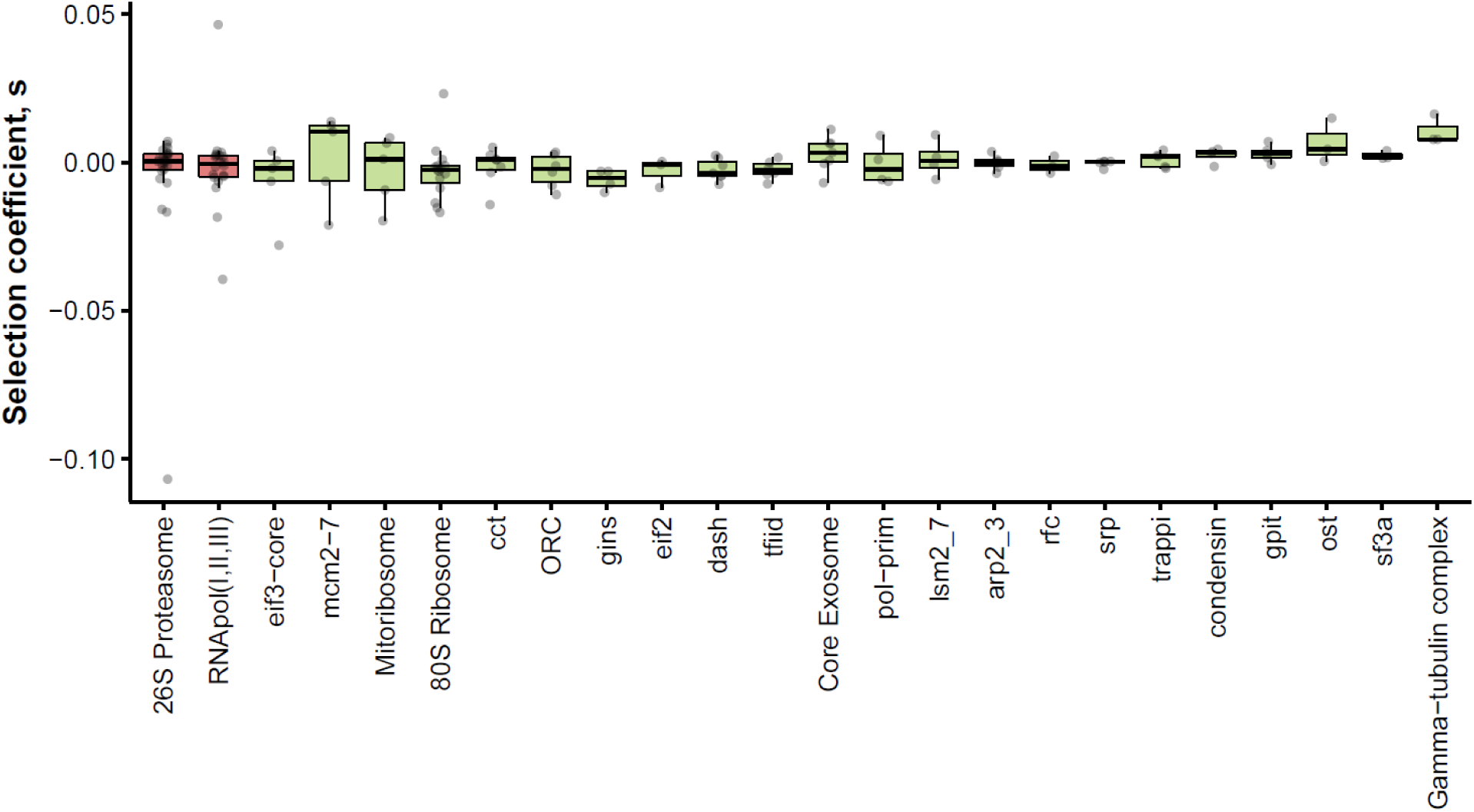
Selection coefficients of strains carrying extra copies of subunits of complexes. The list of proteins that are members of protein complexes was obtained from the Complex Portal Database (https://www.ebi.ac.uk/complexportal/home). Complexes are ordered according to their most deleterious duplication. In red we show the complexes selected for further experiments.

**Figure S6.**
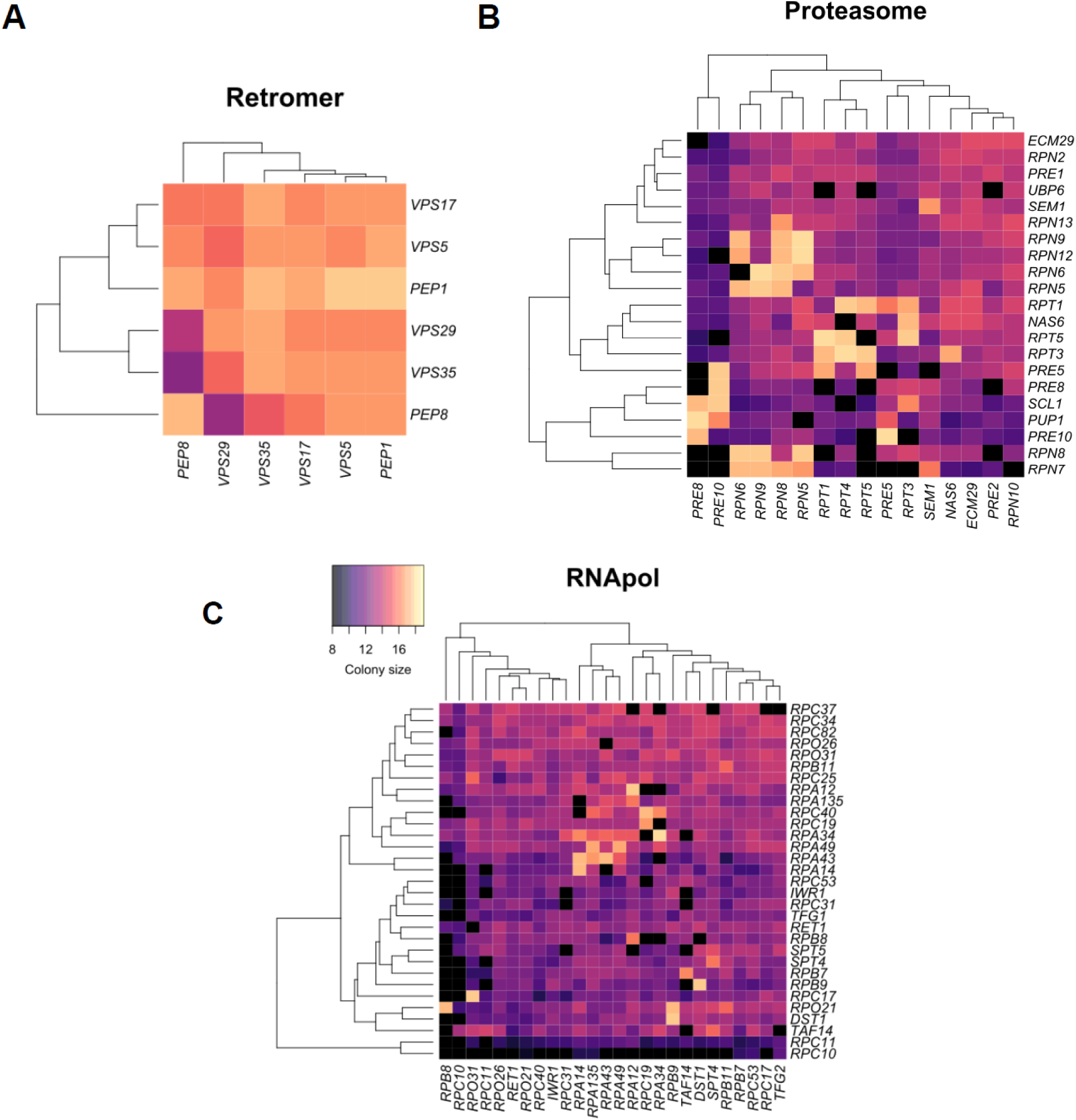
Heatmaps of wild-type interactions of the (**A**) retromer complex; (**B**) 26S Proteasome Complex and (**C**) RNA polymerase I, II, and III. Each tile represents a different diploid strain obtained from the mating of the preys strains (MATα prey-DHFR-F[3]) indicated in the columns and the baits (MATa bait-DHFR-F[1,2]) indicated in the rows. The values on the color key represent the size of the colony of that diploid strain on the fourth day of the MTX II selection plate. White tiles represent missing values. Colony sizes above 14.7 are considered positive interactions (see methods).

**Figure S7.**
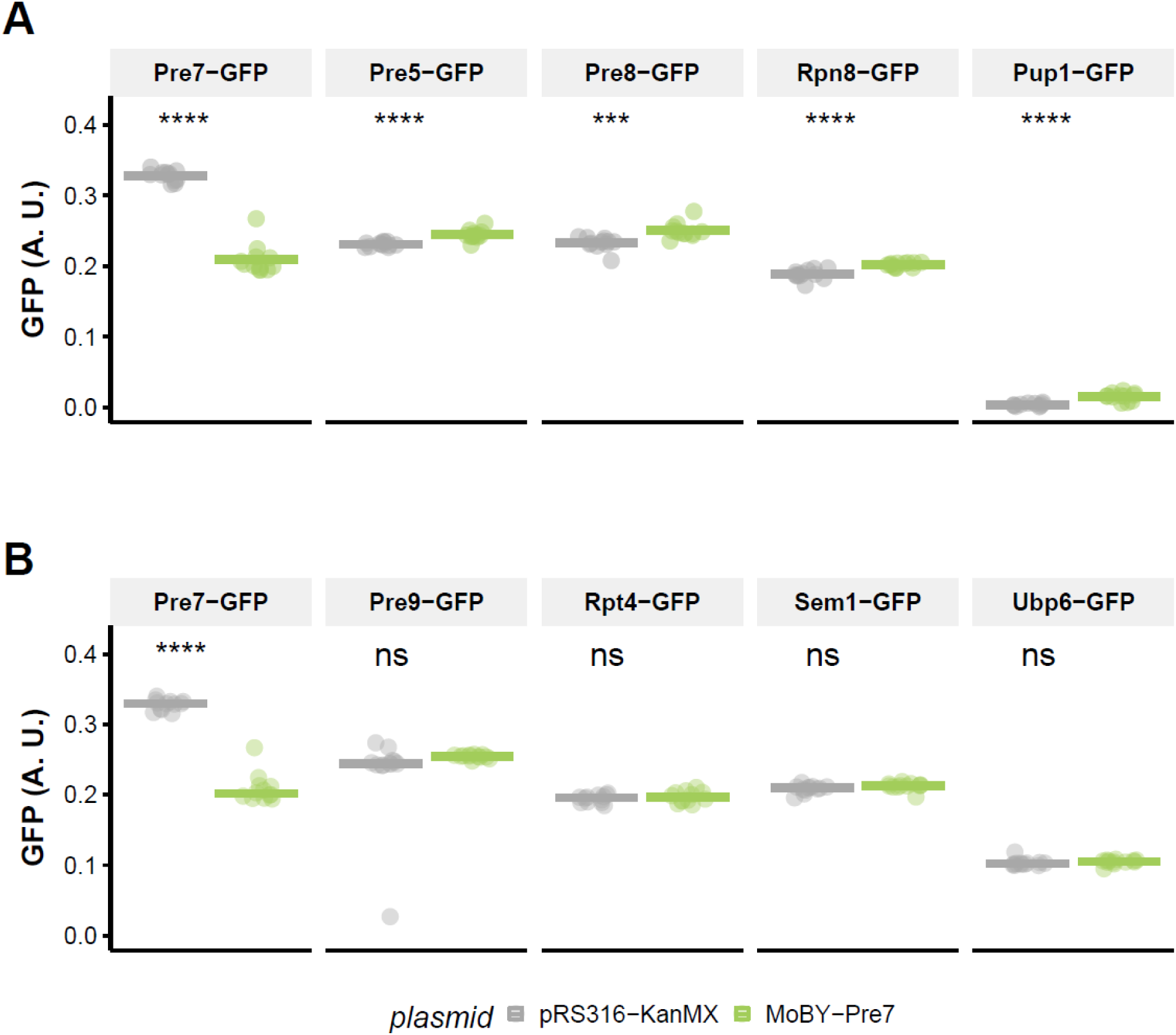
Protein abundance changes of various proteins after *PRE7* duplication. Average GFP signal of proteasome subunits with PPIs (**A**) perturbed and (**B**) non-perturbed by PRE7 duplication. In green, GFP- strains bearing a *PRE7* duplication and, in gray, strains carrying the empty vector as a control. Each dot represents a biological replica and is the mean of up to 5,000 events measured by flow cytometry. The horizontal bar indicates the median of the 12 biological replicas. The GFP signals are normalized with the autofluorescence signal of the parental strain BY4741 not expressing GFP. Each strain was isolated from an independent colony of the transformation plates. On top, we show the *p*-values of Student’s t-test (***: *p* < 0.001; ****: *p* < 0.0001; ns: *p >* 0.05).

**Figure S8.**
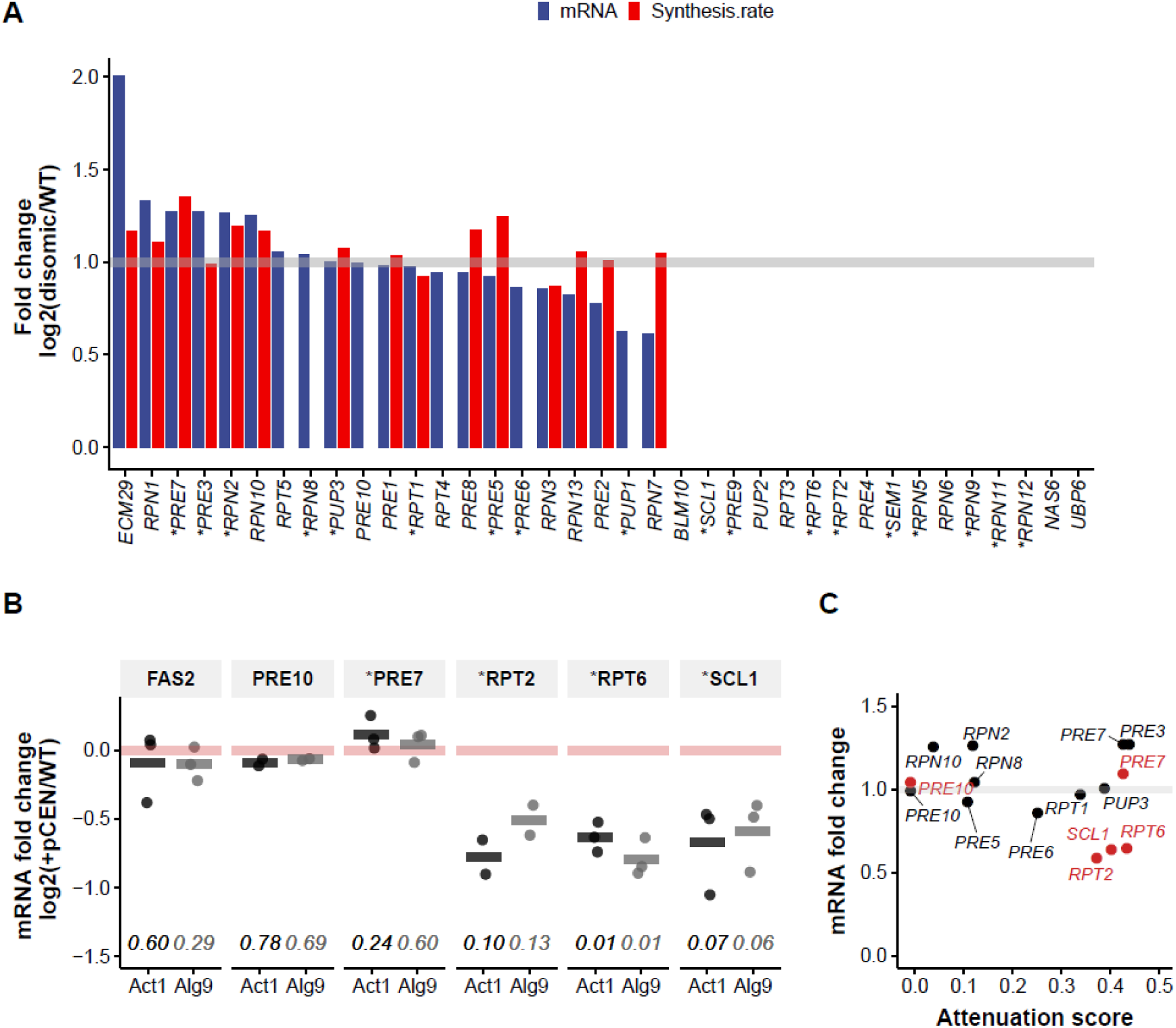
Transcriptomic and translational responses to gene doubling. (**A**) The transcriptional and translational changes in all proteasome subunits reported in disomic chromosomes. Change in mRNA abundance and synthesis rates estimated by ribosome profiling of duplicated proteasome subunits relative to wild-type (single-copy). Blue bars represent mRNA ratios from Dephoure *et al*. (2014). This genome-wide screening estimates the mRNA abundance of individual genes in aneuploid strains relative to wild-type. Red bars represent protein synthesis rate ratios obtained from Taggart *et al*. (2018) and derived from ribosome profiling. The gray line represents the expected value (doubling) when there is no feedback regulation after the duplication of these subunits. (**B**) The mRNA abundance changes in response to duplication. mRNA ratios of xxx-GFP-tagged in strains carrying a pCEN-xxx relative to a WT strain (carrying an empty vector). The red line represents the value expected if there is no change at the mRNA level since we measured only the GFP-tagged transcripts. On the x-axis we indicate the gene transcript used as a reference for the normalization of each mRNA quantification, either Act1 or Alg9, which both give similar results. On the bottom, we show p-values of one-sample Student’s t-test using a true mean of 0. *PRE10* and *PRE7* show similar responses with the plasmid-based duplication and disomic chromosomes shown above. The disomic expression data is, therefore, consistent with the plasmid-based duplication data. *Dephoure et al. did not measure RPT6, SCL1 and RPT2*. We used Fas2 as control outside of the proteasome. In (**A**) and (**B**) we indicate with an asterisk genes that we detected as attenuated after duplication (attenuation score > 0.1, Figure 3). (**C**) Relationship between the mRNA fold change in disomic (black, Dephoure *et al*.) or pCEN(red, this work) strains and attenuation score. While most proteasome subunits are regulated posttranscriptionally, our new data suggest that regulation of *RPT6, SCL1* and *RPT2* is transcriptional.

**Figure S9.**
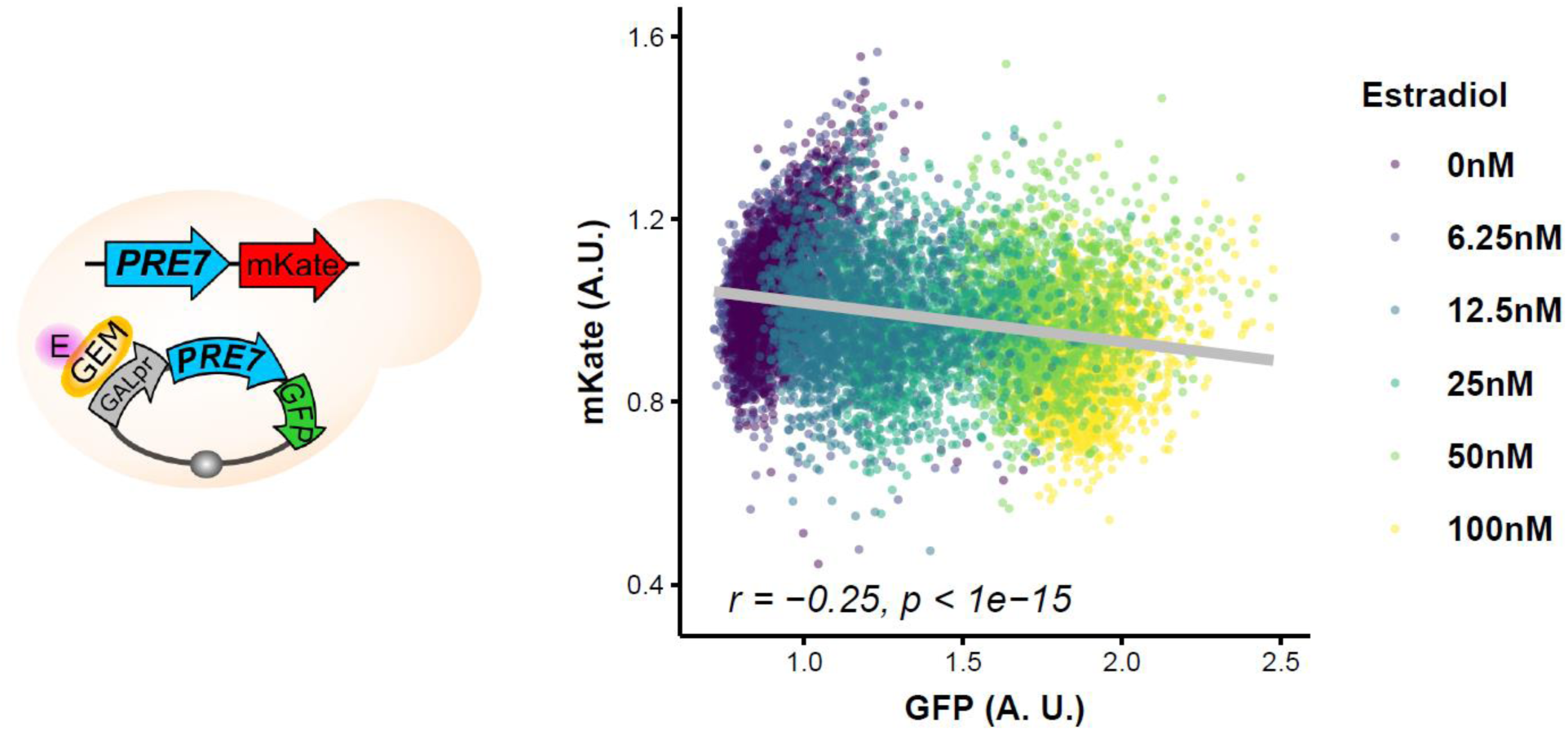
Pre7p attenuation is dosage-dependent. On the left, a cartoon of the strain used for this experiment. The genomic copy of *PRE7* is tagged with mKate. The strain was transformed with a plasmid bearing an extra copy of *PRE7* expressed under the *GAL1* promoter and tagged with GFP. The parental strain that expresses the GEM (yellow oval) chimeric transcriptional regulator (Aranda-Díaz et al. 2017) is activated in the presence of estradiol (pink circle). On the right, the mKate fluorescent signal (*PRE7* chromosome-coded copy) as a function of the GFP signal (plasmid-coded copy). Each point represents an event detected by flow cytometry. We captured up to 5,000 events for each concentration of estradiol. Colors represent different estradiol concentrations (*Supplementary TableS8*).

**Figure S10.**
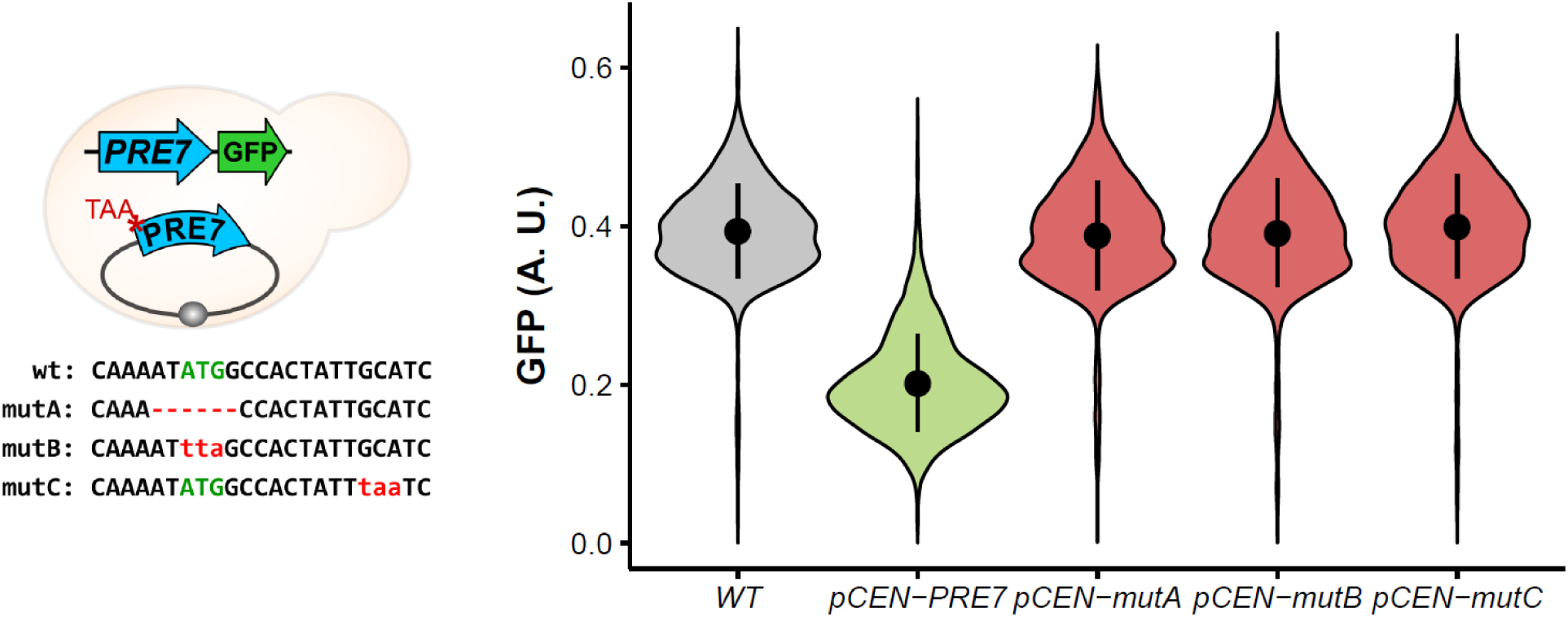
Duplicated *PRE7* copies mutagenized with premature stop codons are not attenuated. On the left, we show a cartoon of the strain bearing a mutagenized *pCEN-PRE7* plasmid and the GFP-tagged *PRE7* genomic copy. In the plasmid *pCEN-mutA*, we deleted the start codon of *PRE7*, on the *pCEN-mutB* plasmid, we replaced the start codon with a stop codon, and in the *pCEN-mutC* plasmid, we replaced the fifth codon with a stop codon. All strains are constructed in the *PRE7-GFP* background (see Methods). On the right, each violin-plot shows the distribution of 5,000 cells GFP normalized signal measured by flow cytometry.

### Supplementary Tables

**Table S1**. Selection coefficients in fluorometry in SC -ura and in 1.2M NaCl.

**Table S2**. Selection coefficients of competition assays performed in cytometry.

**Table S3**. Sensitivity to different z-score thresholds.

**Table S4**. List of preys, baits, and plasmids used for the PCA experiments.

**Table S5**. DHFR-PCA data: colony sizes, *q-values*, and *ps*.

**Table S6**. Cytometry GFP measurements and attenuation scores.

**Table S7**. Cytometry data for duplication of PRE7 in all the other subunits GFP-fusion strains.

**Table S8**. Cytometry GFP data of the tunable promoter system experiments with PRE7.

**Table S9**. Oligonucleotides used in this study (confirmations, GFP-tagging, plasmid construction).

**Table S10**. Media and solutions used in this study.

